# Neural instructive signals for associative cerebellar learning

**DOI:** 10.1101/2022.04.18.488634

**Authors:** N. Tatiana Silva, Jorge Ramírez-Buriticá, Dominique L. Pritchett, Megan R. Carey

## Abstract

Supervised learning depends on instructive signals that shape the output of neural circuits to support learned changes in behavior. Climbing fiber inputs to the cerebellar cortex represent one of the strongest candidates in the vertebrate brain for conveying neural instructive signals. However, recent studies have shown that Purkinje cell stimulation can also drive cerebellar learning, and the relative importance of these two neuron types in providing instructive signals for cerebellum-dependent behaviors remains unresolved. Here we used cell-type specific perturbations of climbing fibers, Purkinje cells, and other cerebellar circuit elements to systematically evaluate their contributions to delay eyeblink conditioning. Our findings reveal that while optogenetic stimulation of either climbing fibers or Purkinje cells are capable of driving learning under some conditions, even subtle reductions in climbing fiber signaling completely block learning to natural conditioning stimuli. We conclude that climbing fibers and corresponding Purkinje cell complex-spike events provide essential instructive signals for associative cerebellar learning.

## INTRODUCTION

Instructive signals are a core component of supervised learning systems. In the brain they are thought to be conveyed by specific classes of neurons that trigger modification of neural pathways that control behavior. Climbing fiber (CF) projections from the inferior olive to the cerebellar cortex have long-been hypothesized to convey neural instructive error signals for various forms of learning, including associative eyeblink conditioning and several forms of motor adaptation^1–9^.

According to the climbing fiber hypothesis, climbing fiber activity drives associative plasticity at parallel fiber inputs to cerebellar Purkinje cells, which forms the neural substrate for learning. There are several lines of evidence in support of this hypothesis. In contrast to typical ‘simple spikes’ (SSpk), which are driven by excitatory parallel fiber inputs, climbing fibers evoke powerful ‘complex spikes’ (CSpk) in cerebellar Purkinje cells (Fig. 1a-d). Complex spikes have a unique electrophysiological signature revealing multiple ‘spikelets’ (Fig. 1d). They are associated with elevations in dendritic calcium and drive heterosynaptic plasticity at parallel fiber-to-Purkinje cell synapses^10–14^. Complex spike activity is associated with sensorimotor errors for a range of behavioral tasks, with the probability of a complex spike often changing in predictable ways with the development of learning^15–22^, and its extinction^23–25^. Moreover, electrical stimulation of climbing fiber pathways is sufficient to substitute for an airpuff unconditioned stimulus (US) to drive eyeblink conditioning in rabbits^26,27^, and recent experiments have shown that optogenetic climbing fiber stimulation can trigger adaptation of the vestibulo-ocular reflex^28,29^ (VOR), while inhibition of climbing fibers can drive extinction of eyeblink conditioning^24,25^.

**Fig. 1.**
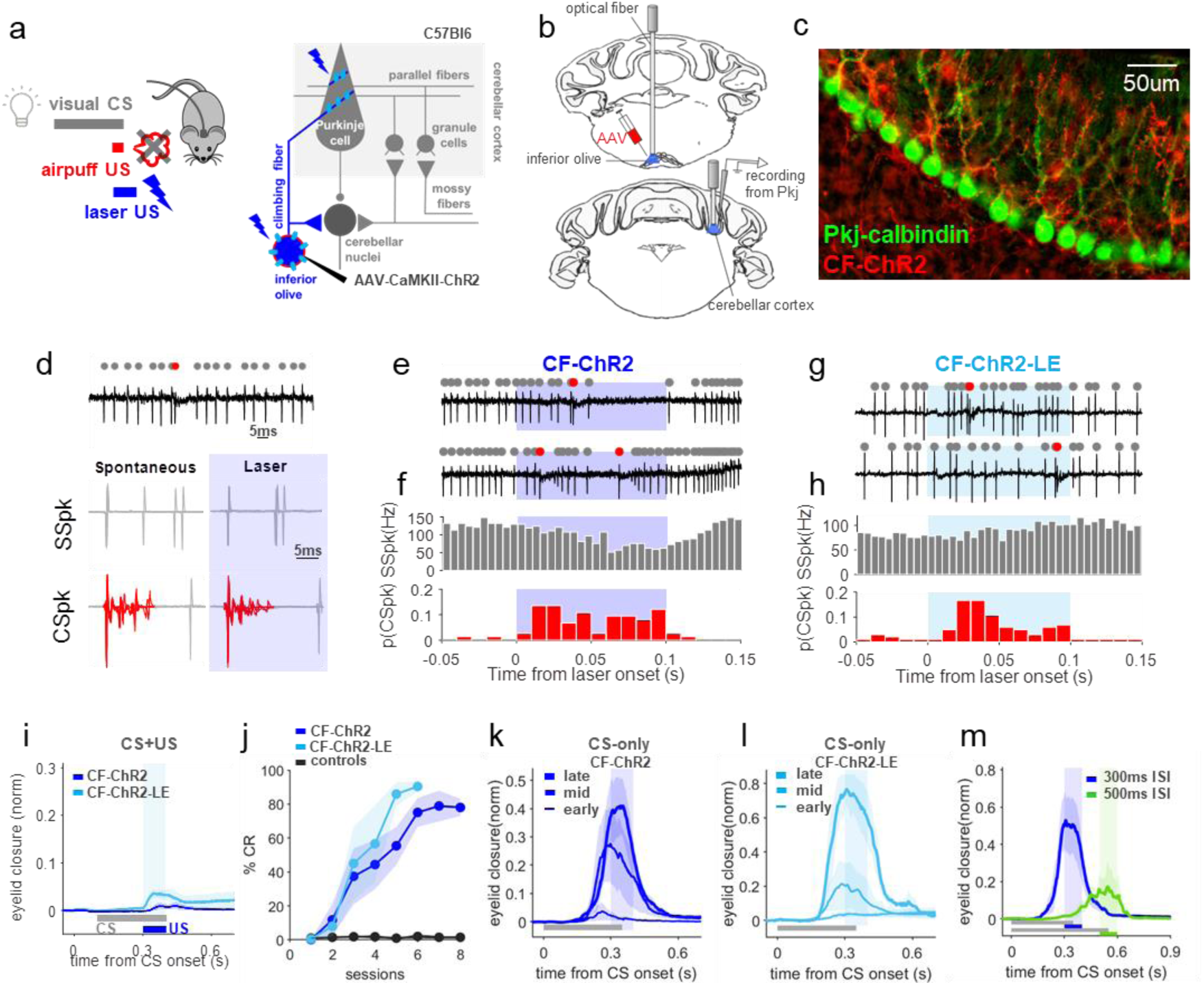
Optogenetic climbing fiber stimulation instructs eyeblink conditioning. **a**, Left, experimental scheme of delay eyeblink conditioning to an optogenetic US. The traditional airpuff-US is replaced by laser stimulation and paired with a visual CS. Right, cerebellar circuit underlying eyeblink conditioning and experimental strategy. Wild-type animals were injected with an AAV-CamKII-ChR2 in the IO and differential sites of photoactivation were used to photoactivate climbing fiber somas in the inferior olive (IO) or terminals in the cerebellar cortex, that project onto Purkinje cells (which also receive input from granule cells/ parallel fibers). **b**, Optical fibers were implanted in either the left IO or the eyelid region of the right cerebellar cortex, where single-unit recordings were performed from Purkinje cells. **c**, Example sagittal section of cerebellar cortex. ChR2 (red) is expressed in climbing fiber inputs to Purkinje cells (green). (Also see Supp. Fig. 1) **d**, Waveforms of simple spikes (SSpk) and complex spikes (CSpk) from an example Purkinje cell during spontaneous and laser epochs. **e**, Two example electrophysiological traces from a Purkinje cell with identified SSpks (grey dots) and CSpks (red) in response to CF-ChR2 laser stimulation in the IO (CF-ChR2-IO, blue); also showing pre- and post-laser epochs. **f**, Population histogram of SSpk firing rate (grey) and CSpk probability (red) (n=74 trials, N=4 units from 2 animals). CSpks: spont. vs laser, P=0.02*, Student’s paired t-Test; SSpks: spont. vs laser, P=0.22n.s., Student’s paired t-Test. **g,h** same as *e,f* but for CF-ChR2 low expression animals (CF-ChR2-LE-IO; n=102 trials, N=8 cells from 2 animals). CSpks: spont. vs laser, P=0.01*, Student’s paired t-Test; SSpks: spont. vs laser, P=0.11n.s., Student’s paired t-Test. **i**, Average eyelid closure trace from CS+US trials of the first training session show no reflexive eyeblink to CF-ChR2-IO laser stimulation (N=7 mice, blue) and a very small eye twitch to laser stimulation in CF-ChR2-LE-IO animals (N=4 mice, light blue). **j**, Percentage of trials in which conditioned responses were observed (%CR) across daily training sessions of animals trained with a CF-ChR2-IO (N=7 mice, blue) or CF-ChR2-LE-IO laser US (N=4 mice, light blue). Controls: wildtype mice (no ChR2 expression) with a fiber implanted in the IO presented with IO-laser US (N=2 mice, black); %CR at last learning session: CF-ChR2-IO vs controls, P= 1.7726e-04***, Student’s t-Test; CF-ChR2-LE-IO vs controls, P=4.0836e-05***, Student’s t-Test; CF-ChR2-IO vs CF-ChR2-LE-IO, P=0.115n.s., Student’s t-Test. **k**, Average eyelid closure traces from CS-only trials of training sessions 2, 4 and 8 for the experiments of CF-ChR2-IO shown in *j*. Shaded rectangle indicates where in the trial the US would have appeared. **l**, same as k but for training sessions 2, 4 and 6 of CF-ChR2-LE-IO. **m**, Average eyelid closures from CS-only trials after training to a CS+US interstimulus interval (ISI) of 300ms (blue, N=4 mice), and 500ms (green, N=4 mice). Peak eyelid closure time on last session of learning: 300ms vs 500ms ISI, P=0.01*, Student’s paired t-Test.

However, significant confusion and controversy remain, particularly regarding the necessity of climbing fiber instructive signals and complex spike-driven plasticity for learning^8,30–35^. For instance, there is substantial experimental support for an alternative model which posits that Purkinje cell simple spike modulation, rather than climbing fiber-driven complex spikes, could provide relevant instructive signals for learning^4,36^. This hypothesis stems from the observation that sensorimotor errors that drive climbing fiber activity and subsequent Purkinje cell complex spikes are often tightly linked to rapid reflexive, corrective movements. Crucially, Purkinje cell simple spike output often correlates with these corrective movements^34,37,38^, raising the possibility that they could provide their own instructive signals for plasticity – either in addition to, or independently of, complex spike activity^30,36,39–44^. In particular, Purkinje cell simple spike modulation could instruct plasticity in the downstream cerebellar nuclei^5,36^, an idea that also has support from in vitro experiments of synaptic plasticity^45,46^.

Seemingly consistent with a possible instructive role for Purkinje cell simple spike modulation, recent work has demonstrated that pairing optogenetic stimulation of Purkinje cells, which effectively modulates simple spike activity, with ongoing movements can drive motor adaptation in multiple systems^28,47,48^. However, it is not clear whether this optogenetically-evoked learning results from modulation of simple spike output, and/or from the generation of complex spike-like dendritic calcium signals in Purkinje cells that instruct plasticity in the cerebellar cortex^48^.

Just as it has not been clear whether Purkinje cell simple spikes could provide alternative instructive signals, it has also not been clear whether climbing fiber signaling is absolutely required for cerebellar learning. Although Purkinje cell CSpk activity often correlates with sensorimotor errors that drive behavioral learning, the extremely low rates of CSpk activity and high proportion of ‘spontaneous’ CSpks that appear not to correspond with identifiable task parameters complicate a definitive interpretation of CSpks as instructive signals^34^. Moreover, much of the evidence to date that has been interpreted as supporting a causal role for climbing fiber instructive signals for cerebellar learning has come from lesion studies^49^, pharmacological inactivations^24,31^, and electrical perturbations^50,51^ of the inferior olive. These manipulations lack both cell-type and temporal specificity and are likely to have substantial additional, unintended effects on the olivocerebellar circuit^52^ that are extremely difficult to control for. Until now, there has not been a precise way to selectively perturb evoked climbing fiber activity while leaving olivocerebellar function otherwise intact.

Here we used cell-type specific perturbations of climbing fibers, Purkinje cells, and other circuit elements to test their sufficiency and necessity as instructive signals for associative cerebellar learning. We combined behavioral, optogenetic and electrophysiological approaches to dissociate climbing fiber inputs and complex spike activity from reflexive movements and simple spike modulation. We find that optogenetically-evoked complex spikes can substitute for an airpuff unconditioned stimulus to induce learning, even in the absence of an evoked blink, while temporally precise optogenetic silencing of climbing fibers completely blocks learning. Direct optogenetic stimulation of Purkinje cells can also drive learning; however, this effect was dissociable from both simple spike modulation and the corresponding evoked blink. Finally, simple ChR2 expression in climbing fibers is associated with a subtle decrease in Purkinje cell complex-spike probability that abolishes learning to a sensory unconditioned stimulus. Together, our results support a necessary and sufficient role for climbing fibers and corresponding Purkinje cell complex spike-events as instructive signals for associative cerebellar learning.

## RESULTS

We investigated neural instructive signals for delay eyeblink conditioning in head-fixed mice walking on a motorized treadmill^53,54^. In classical eyeblink conditioning experiments (Fig. 1a), a neutral conditioned stimulus (CS, here a white light LED) is paired with an unconditioned stimulus (US, usually a puff of air directed at the eye) that reliably elicits an eyeblink unconditioned response (UR) and serves as an instructive signal for learning. Climbing fibers from the dorsal accessory part of the inferior olive respond to the airpuff-US and project to the contralateral cerebellum, where they drive ‘complex spikes’ (CSpk) in Purkinje cells in the cerebellar cortex (Fig. 1a,b). Information about the CS is conveyed to the cerebellum by mossy fibers that synapse onto granule cells, whose axons form parallel fiber inputs that modulate ‘simple spikes’ (SSpk) in Purkinje cells. Pauses in simple spike activity in the eyelid region of cerebellar cortex are associated with eyelid closures^55–58^. Here we use genetic circuit dissection to distinguish between competing models in which CSpks and/or SSpk modulation provide instructive signals for eyeblink conditioning. We first asked whether direct optogenetic stimulation of climbing fibers could substitute for a sensory (airpuff) US to drive behavioral learning (Fig. 1).

### Optogenetic climbing fiber stimulation is sufficient to drive eyeblink conditioning

To specifically target climbing fibers (CF), we injected a virus that allows for expression of ChR2 under control of the CaMKIIα promoter^28^ (AAV-CaMKII-ChR2; here termed CF-ChR2) into the dorsal accessory inferior olive (IO) of wildtype mice (Fig. 1a,b; Methods). With this strategy we observed selective labeling of neurons in the IO and climbing fibers in the cerebellar cortex (Fig. 1c; Supp. Fig. 1a-f). An optical fiber was placed either in the dorsal accessory IO^25,26,28^ (CF-ChR2-IO), targeting cell bodies of eyeblink-related climbing fibers, or in the eyelid region of the cerebellar cortex^56–59^, targeting CF terminals (CF-ChR2-Ctx; Fig. 1b; Supp. Fig. 1; Methods). Laser stimulation at both sites evoked robust postsynaptic complex spike responses in cerebellar Purkinje cells, with waveforms matching those of spontaneous complex spikes (Fig. 1d-f; Supp. Fig. 1g,h). Similar electrophysiological responses were observed in mice expressing ChR2 at standard levels, or with a slightly reduced viral titer (CF-ChR2-LE; 5-fold lower titer; Fig. 1g,h).

To test the sufficiency of climbing fiber activity for the acquisition of learned eyelid responses, we paired a neutral visual CS with optogenetic CF stimulation in the absence of any sensory US (CF-ChR2-US; Fig. 1a). Laser stimulation alone did not elicit robust eyelid closures (Fig. 1i). Despite the absence of an eyeblink unconditioned response (UR) to the optogenetic-US, conditioned eyelid closure responses (CRs) gradually emerged in response to the visual CS following repeated CS+US pairing (Fig. 1j,k). Similar learning was observed with both expression levels (Fig. 1j-l) and for fiber placements either in the IO (CF-ChR2-IO; Fig. 1d-l) or the cerebellar cortex (CF-ChR2-Ctx; Supp. Fig. 1g-k; though note the subtly different CR and CSpk timings in the two cases). Moreover, learning was also observed in separate experiments in which we targeted ChR2 expression to glutamatergic IO neurons with a transgenic, rather than viral strategy, by crossing vGlut2-Cre mice with ChR2-floxed mice^60^ (*vglut2-Cre;ChR2*; Methods) and placing the fiber in the IO (vGlut2-ChR2-IO; Supp. Fig. 1m-p).

In general, the properties of learning to an optogenetic CF-US matched those of normal sensory CS+US conditioning in wildtype mice^53,54^. Learning to an optogenetic-US was unilateral (specific to the eye contralateral to the IO and ipsilateral to the corresponding cerebellar cortex; Supp. Fig. 1l) and emerged over several days, with both frequency and amplitude of learned eyelid closures increasing gradually across sessions (Fig. 1j-l, Supp. Fig.1i,o,p; %CR at last learning session: CF-ChR2-IO vs airpuff-US controls in Supp. 1o, P=0.29n.s., Student’s t-Test; CF-ChR2-LE-IO vs airpuff-US controls in Supp. 1o, P=0.54n.s., Student’s t-Test). Learning also extinguished appropriately upon cessation of CS+US pairing, when CSs were presented alone (Supp. Fig. 1i).

A central feature of eyeblink conditioning is the appropriate timing of the CR, so that its peak generally coincides with the expected time of the arrival of the US^61,62^. This appropriate timing was also observed for learning to an optogenetic US (Fig. 1k,l; Supp. Fig. 1j,k,p). Moreover, when the interval between CS and CF-ChR2-US onset was shifted from 300ms to 500ms, mice adapted the timing of their learned responses^54,62^ (Fig. 1m).

These results indicate that optogenetic climbing fiber activation is sufficient to substitute for an airpuff-US to instruct delay eyeblink conditioning.

### Optogenetic stimulation of Purkinje cells can substitute for a US to drive learning

Optogenetic stimulation of Purkinje cells has previously been shown to instruct motor adaptation of limb and eye movements^28,47,48^. To investigate whether this was also true for delay eyeblink conditioning, we placed an optical fiber at the eyelid region of the cerebellar cortex of transgenic mice expressing ChR2 under the L7 Purkinje-cell specific promoter (*L7-Cre;Chr2* mice; Fig. 2a,b). Consistent with previous studies^47,53,63–68^, in vivo electrophysiological recordings confirmed an increase in Purkinje cell simple spike activity at the onset of low-medium intensity optogenetic stimulation, followed by a slow decrease below the baseline firing rate upon the cessation of stimulation, without significant changes in complex spike activity (Fig. 2c,d, Supp. Fig. 2d).

**Fig. 2.**
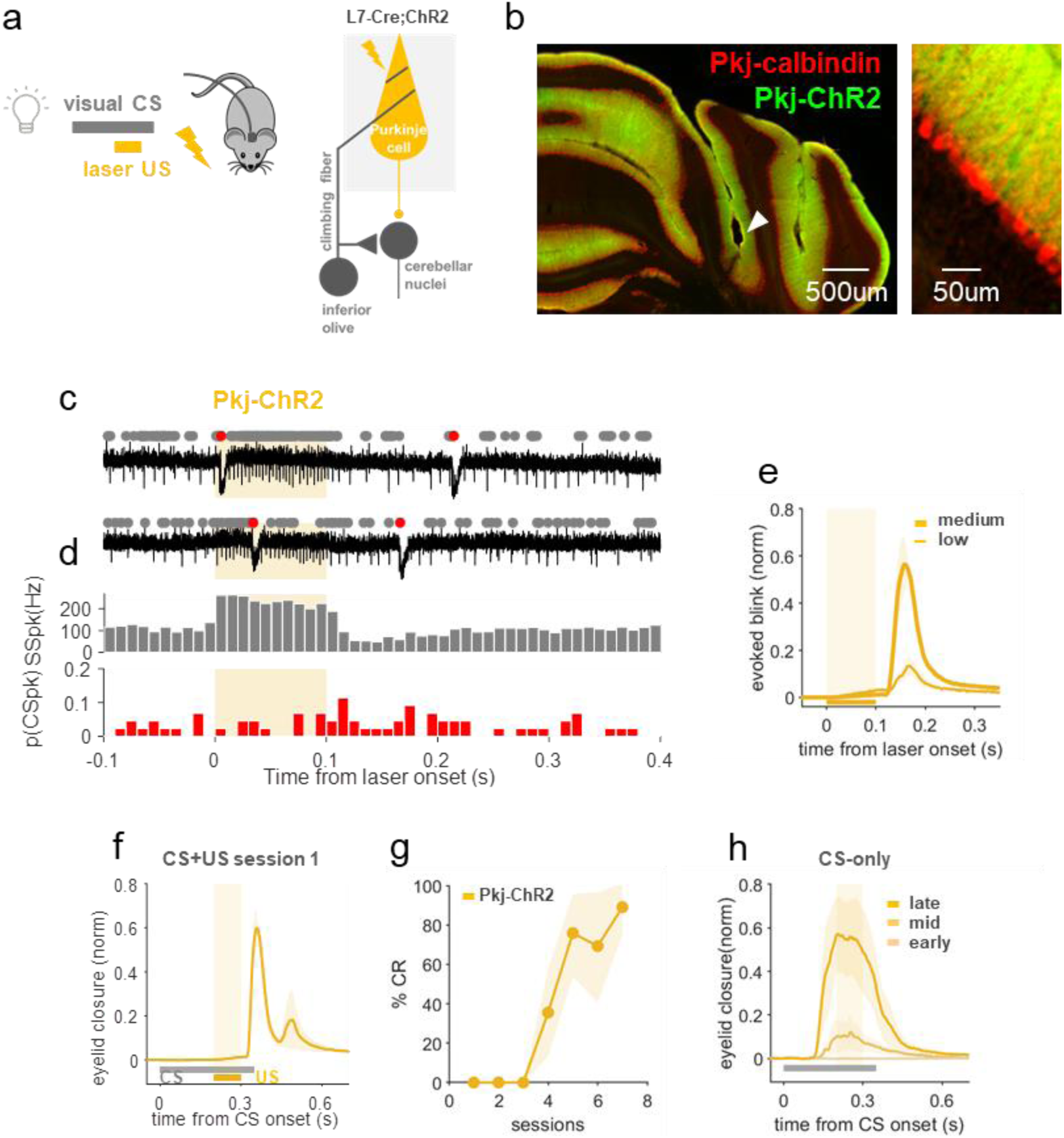
Optogenetic stimulation of Purkinje cells can substitute for a US to drive learning. **a**, Experimental scheme. L7-Cre;ChR2 mice were used to photostimulate Purkinje cells, which served as a US for conditioning. **b**, Example coronal section of cerebellar cortex indicating fiber placement in the eyelid area of the cerebellar cortex (white arrow) and labeling Purkinje cell ChR2 expression (green) and calbindin (red). **c**, Example electrophysiological traces of Purkinje cell SSpks (grey dots) and CSpks (red) in response to Pkj-ChR2 laser stimulation (orange shading). **d**, Population histogram of SSpk rate (grey) and CSpk probability (red; n=44 trials, N=2 units from 2 mice); see Supp. Fig. 2d for stats. **e**, Average eyelid closures evoked by low and medium-power Pkj-ChR2 stimulation (N=4 mice). Note the blink at stimulus offset. Peak amplitude of evoked blink: low vs medium power, P=0.047*, Student’s paired t-Test. **f**, Average eyelid closures on CS+US trials in the first training session showing the blink evoked by Pkj-ChR2-US laser stimulation (N=4 mice). **g**, %CR across training sessions to a Pkj-ChR2-US (N=4 mice, plotted as in Fig. 1j). **h**, Average eyelid traces from CS-only trials of sessions 2, 4, and 7 of the experiments in *g*.

The post-stimulation inhibition of Purkinje cell simple spikes is consistently observed upon optogenetic stimulation *in vivo*^47,63–68^, but not *in vitro* with synaptic activity blocked^69,70^, and likely reflects synaptically-mediated network effects^71–73^. In the eyelid region of cerebellar cortex, eyelid closures are associated with decreases in Purkinje cell simple spike firing rate^55–58^. Consistent with this, and with previous optogenetic studies^53,56^, we found that optogenetic Purkinje cell stimulation with low-medium laser intensities resulted in eyelid closures upon stimulus offset, and that their amplitudes scaled as a function of laser intensity (Fig. 2e).

When a visual CS was consistently paired with a US consisting of optogenetic stimulation of Purkinje cells that drove increased simple spike activity at onset and a blink at laser offset (Pkj-ChR2; Fig. 2f), robust conditioned responses gradually emerged (Fig. 2g,h). Rates of learning and conditioned response amplitudes were comparable to those obtained with an airpuff-US^53^ (%CR at last learning session: Pkj-ChR2 vs airpuff-US controls in Supp. 1o, P=0.92n.s., Student’s t-Test) and also with the CF-ChR2-US used in Fig. 1j (%CR at last learning session: Pkj-ChR2 vs CF-ChR2-IO, P=0.24n.s., Student’s t-Test; Pkj-ChR2 vs CF-ChR2-LE-IO, P=0.75n.s., Student’s t-Test). Notably, conditioned responses were timed so that the peak eyelid closure coincided with the expected time of the onset of optogenetic stimulation (Fig. 2h).

The results of Fig. 2 suggest that direct optogenetic perturbation of Purkinje cells can substitute for an airpuff unconditioned stimulus to act as instructive signals to drive eyeblink conditioning. However, they do not allow us to disentangle possible contributions of increases and/or decreases in Purkinje cell simple spikes or evoked eyelid closures. In the next set of experiments we systematically altered the temporal relationships between these candidate instructive signals by varying laser timing, duration, and intensity.

### Onset of Purkinje cell optogenetic stimulation drives learning independently of simple spike modulation or an evoked blink

The well-timed conditioned responses observed in eyeblink conditioning are thought to be a consequence of plasticity mechanisms acting within the cerebellar cortex that associate postsynaptic calcium events (usually complex spikes) in Purkinje cells with a particular set of parallel fiber inputs active within a particular temporal window from the onset of the conditioned stimulus^4,6,7,13,61,74^. We first asked whether learning to an optogenetic Pkj-ChR2 US would yield well-timed conditioned responses to different CS-US intervals (Fig. 3b,c). Indeed, extending the interstimulus interval between CS and Pkj-ChR2-US onset from 200ms to 400ms revealed appropriate corresponding shifts in CR timing (Fig. 3d-e).

**Fig. 3.**
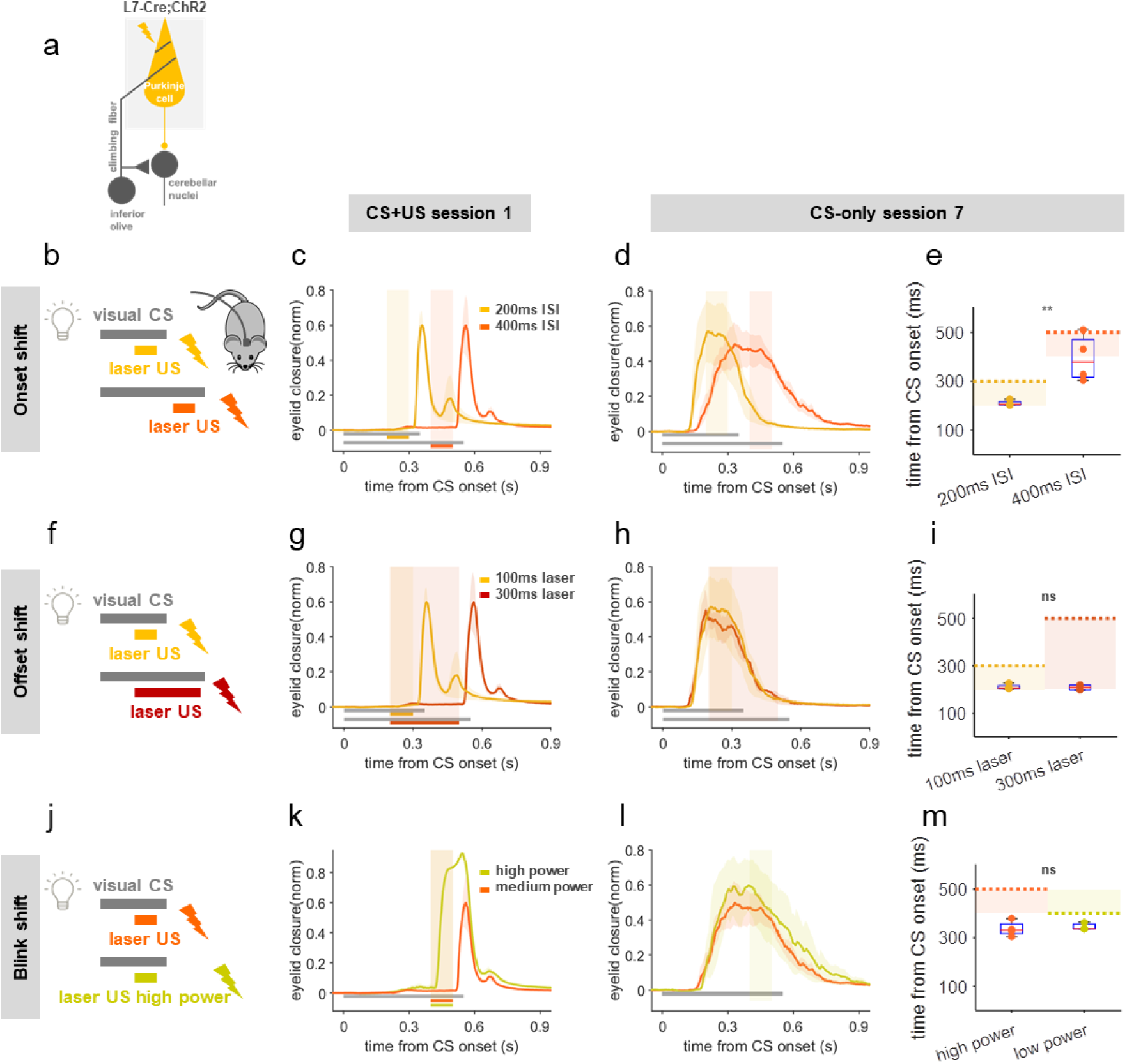
Learning evoked by optogenetic Purkinje cell stimulation is temporally coupled to stimulation onset, and not evoked blinks or simple spike modulation. **a**,**b**,**f**,**j**, Schemes for Pkj-ChR2-US experiments in which stimulation onset timing, duration, and intensity were varied systematically to dissociate candidate instructive signals (also see Supp. Fig. 2). **b**, US onset was shifted to obtain CS+US interstimulus intervals of 200 (yellow) or 400ms (orange). **c**, Before training, evoked blinks on CS+US trials occurred at US offset in both conditions (N=4 mice each). **d**, Following training, the temporal profile of eyelid closures on CS-only trials depended on the timing of US-onset. **e**, The timing of peak eyelid closures on CS-only trials was later for the longer ISI. Peak eyelid closure time: 200ms vs 400ms ISI, P=0.009**, Student’s t-Test. Shaded rectangles indicate laser US duration and dashed lines indicate blink onset. Each dot is one mouse and box plots indicate 25^th^-75^th^ percentiles with whiskers extending to most extreme data points. **f**, US duration was adjusted so that CS+US onset times were identical, but US offset (and blink) timing varied with respect to the CS. **g**, US-evoked blinks on CS+US trials occurred at stimulus offset (note the temporal correspondence with the blinks in *c*). Also see Supp. Fig. 2b-e. **h**,**i** the temporal profiles of CRs did not depend on the timing of stimulus offset or the evoked blink (peak eyelid closure time: 100ms vs 300ms laser duration, P=0.87n.s., Student’s t-Test), but rather, stimulation onset. **j**,**k,** Laser intensity was adjusted to evoke a blink (associated with a decrease in simple spikes, see Supp. Fig. 2f-h) either at laser offset (orange, as above), or, with higher intensities, at laser onset (lime green; N=3 mice). Laser-US timings and durations were identical in the two conditions. **l**,**m** CR temporal profile depended only on the time of stimulation onset and did not vary with the timing of the evoked blink (peak eyelid closure time: high vs medium laser power, P=0.67n.s., Student’s t-Test) or the direction of simple spike modulation (see Supp. Fig. 2f-h).

Having thus established that learned responses to an optogenetic Pkj-US can be appropriately timed, we next varied the duration of laser stimulation (Fig. 3f; Supp. Fig. 2b-e) to determine whether conditioned responses were timed to match the increase in simple spike activity at the onset of Pkj-ChR2-stimulation, or its offset and the associated blink. To do this we compared conditions in which the onset of Purkinje cell stimulation was presented at the same interstimulus interval relative to the CS, but the offset (and its respective blink) differed by 200ms (Fig. 3f,g; Supp. Fig. 2b-e). The conditioned response amplitudes and timings were identical in the two groups (Fig. 3h,i). This result suggests that events associated with the *onset* of optogenetic stimulation, and not the decrease in simple spike rate or the corresponding blink evoked at laser offset, are crucial for learning driven by optogenetic stimulation of Purkinje cells.

We next exploited the relationship between laser power, simple spike modulation, and timing of the evoked blink to further disambiguate which consequences of the onset of optogenetic stimulation were responsible for optogenetically-driven learning. At higher powers, laser stimulation induces a pause in Purkinje cell simple spike activity at laser onset (Supp. Fig. 2f-h), likely due to depolarization block^75,76^. Consistent with this and the well-established relationship between Purkinje cell simple spike inhibition and eyelid closures, we found that increasing laser intensity also led to a temporal shift in the timing of the optogenetically-evoked blink – from laser offset, to laser onset (Supp. Fig. 2j). We took advantage of this feature to compare learning under conditions in which the timing and duration of the optogenetic ‘US’ stimulation were identical, but laser power was adjusted to invert the direction of simple spike modulation and shift the timing of the evoked blink from laser offset to laser onset (Fig. 3j,k). As in the experiment presented in Fig. 3f-i, here, too, we found that conditioned response timing depended only on the timing of laser onset, and not the timing of the evoked blink on the paired trials (Fig. 3l,m). This again suggests that the relevant instructive signal for learning occurs at the onset, and not offset, of Purkinje cell optogenetic stimulation. Moreover, because of the switch from increases in simple spike activity to pauses in simple spike activity evoked by laser onset at high intensities (Supp. Fig. 2f-h), it further dissociates laser onset from the modulation of Purkinje cell simple spikes as the relevant instructive stimulus for learning.

Taken together, the results of Figs. 2 and 3 suggest that while Purkinje cell optogenetic stimulation can substitute for an airpuff-US to drive eyeblink conditioning, the effective instructive stimulus driving this learning is tightly linked to the onset of laser stimulation, but independent of either the direction of simple spike modulation or the blink that it evokes. One possible explanation for this finding would be if Pkj-ChR2 stimulation elevates dendritic calcium, triggering complex spike-like events that are capable of driving learning, as has been recently demonstrated for VOR adaptation^48^. Consistent with this possibility, we observed electrophysiological signatures of complex spike-like events at the onset of Pkj-ChR2 stimulation at higher stimulation intensities (Supp. Fig. 2i).

If Pkj-ChR2-US stimulation drives eyeblink learning through the generation of dendritic complex spike-like events, then we would predict that Purkinje cell SSpk modulation driven by synaptic inputs rather than direct optogenetic stimulation might not be sufficient to induce learning, even if it was strong enough to evoke a blink. To test this prediction, we replaced direct Pkj-ChR2 stimulation with optogenetic stimulation of cerebellar granule cells, whose axons form parallel fiber inputs to Purkinje cells (Gabra6-ChR2; Fig. 4a,b). As we have previously shown^53^, granule cell stimulation drives a blink at laser onset, consistent with net inhibition of Purkinje cells via molecular layer interneurons^77^ (Fig. 4c). Although this stimulation effectively modulated Purkinje cell simple spikes (Fig. 4d,e) and drove a blink (Fig. 4c,f), it did not generate a complex spike-like-event (Fig. 4d,e), and pairing it with a visual conditioned stimulus did not result in learning (Fig. 4 f-h).

**Fig. 4.**
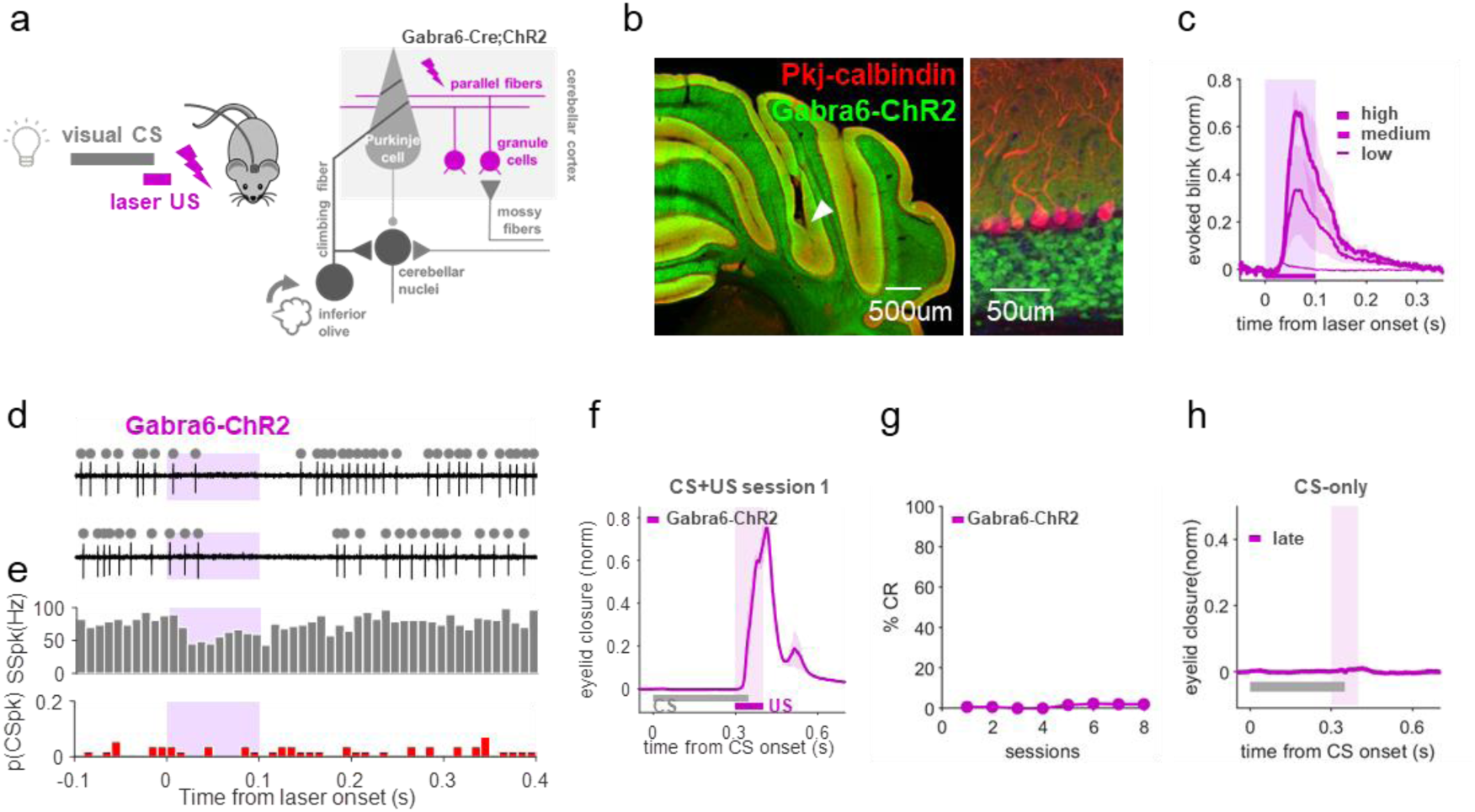
Optogenetic stimulation of cerebellar granule cells drives a blink but not learning. **a**, Experimental scheme. Gabra6-Cre;ChR2 mice were used to photostimulate cerebellar granule cells, which served as a US for conditioning. **b**, Example coronal sections of cerebellar cortex showing expression of ChR2 in granule cells (Gabra6-ChR2, green; Pkj-calbindin, red) and fiber placement in the eyeblink area of the cerebellar cortex (white arrow). **c**, Gabra6-ChR2 laser stimulation evoked intensity-dependent eyelid closures at stimulation onset (N=6 mice). Peak amplitude of evoked blink: medium vs high power, P=0.03*, Student’s paired t-Test. **d**, Example electrophysiological traces of Purkinje cell SSpk (grey dots) and CSpk (red) modulation to Gabra6-ChR2 laser stimulation (purple shading). **e**, Population histograms (n=56 trials, N=3 cells from 2 mice), show decrease in SSpks (spont. vs laser, P=1.28e-05***, Student’s paired t-Test) and no change in CSpks (spont. vs laser, P=1n.s., Student’s paired t-Test) upon laser stimulation. **f**, Average eyelid closures on CS+US trials of the first training session show the blink evoked by Gabra6-ChR2 laser stimulation (purple, N=6 mice). **g**, %CR across sessions (N=6 mice). **h**, Average eyelid traces from CS-only trials of the last training session show no learning (purple, N=6 mice).

The most parsimonious interpretation of the data presented in Figs. 2-4 and Supp. Fig. 2 is that optogenetic stimulation of Purkinje cells drives eyeblink conditioning through cell-autonomous, complex spike-like, events associated with stimulation onset^48^ (Fig. 7).

### Optogenetic inhibition of the inferior olive blocks eyeblink conditioning

We next used several complementary approaches to ask whether climbing fiber activity is required for delay eyeblink conditioning to a sensory airpuff-US. First, to inhibit climbing fibers specifically at the time of US, we injected a virus that allows for expression of the optogenetic inhibitor Jaws^78^ under control of the CaMKIIα promoter (AAV-CaMKII-Jaws) into the inferior olive of wildtype mice (Fig. 5a). We observed selective labeling of neurons in the IO and climbing fibers in the cerebellar cortex (Fig. 5b). As before, an optical fiber was placed in the dorsal accessory IO.

**Fig. 5.**
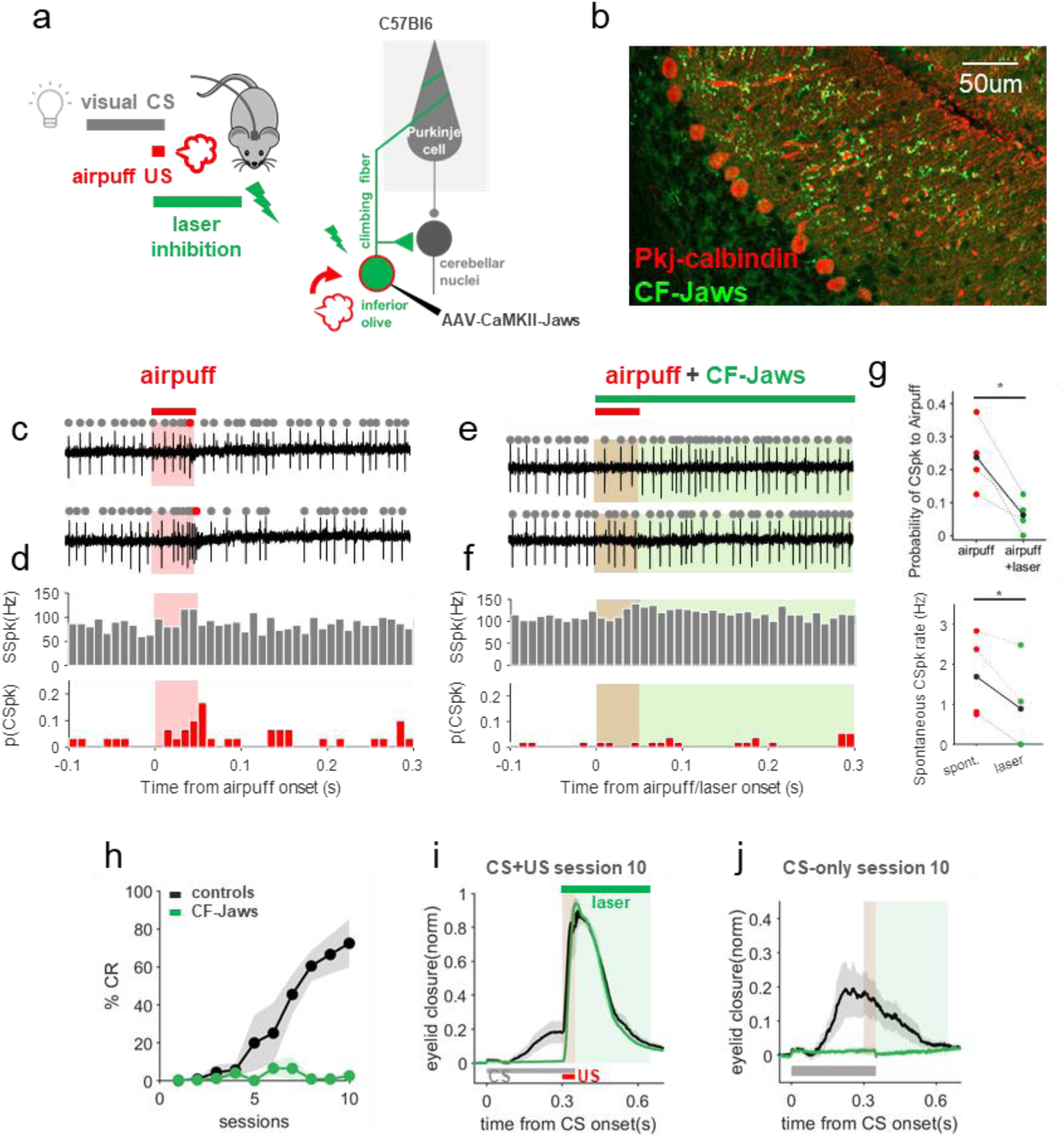
Inhibition of the inferior olive that blocks airpuff-US-driven CSpks eliminates eyeblink conditioning. **a**, Experimental scheme. Photoinhibition of climbing fibers started at airpuff-US onset in CS+US trials. (Duration was randomized to avoid consistently timed rebound excitation, see Methods.) Wild-type animals were injected with AAV-CamKII-Jaws in the inferior olive (IO) where an optical fiber was also placed to photoinhibit climbing fibers. **b**, Example sagittal section of cerebellar cortex. Jaws (green) is expressed in climbing fiber inputs to Purkinje cells (red). **c**, Two example electrophysiological traces from a Purkinje cell with identified SSpks (grey) and CSpks (red) in response to airpuff stimulation (red shading). **d**, Population histogram of SSpks (grey) and CSpks (red) (n=29 trials, N=4 units from 2 animals). **e,f** same as *c,d* but paired with CF-Jaws laser inhibition (green; n=68 trials, same units as in *c,d*). Spontaneous SSpk rate pre- and during-laser epochs (N=4 units; P=0.42n.s., Student’s paired t-Test). **g**, Top, average probability of CSpks in airpuff only vs. airpuff+laser trials (N=4 units; P=0.028*, Student’s paired t-Test). Bottom, spontaneous CSpk rate pre- and during-laser epochs (N=4 units; P=0.026*, Student’s paired t-Test). Each circle represents a unit, linked through conditions by a dotted line; the black solid circles and line represent the average. **h**, %CR across sessions with (green, N=4 mice) and without (black, N=4 mice) CF-Jaws laser inhibition. %CR at last learning session: CF-Jaws vs controls, P= 0.0015**, Student’s t-Test. Control animals expressed Jaws in climbing fibers but no laser was presented. **i**, Average eyelid closure traces from CS+US trials of the last training session of the experiment shown in *h* reveal an absence of learning in the laser inhibition condition despite the intact unconditioned reflex to the airpuff-US. Shaded rectangle indicates where in the trial the US (red) and the laser (green) appeared. **j**, Same as for *i*, but for CS-only trials. Shaded rectangles indicate where US and laser would have appeared.

Photoinhibition of climbing fibers blocked airpuff-driven CSpks (Fig. 5c-g) and also reduced spontaneous complex spiking during laser presentation (Fig. 5g). Moreover, laser inhibition of climbing fibers at the time of the airpuff-US completely prevented learning in CF-Jaws animals (Fig. 5h-j, green), while control mice expressing Jaws in climbing fibers that did not receive laser inhibition learned normally (Fig. 5h-j, black). Notably, the learning impairment could not have been due to an overall inability to respond to the US, as the reflexive blink to the airpuff-US (unconditioned response) was intact (Fig. 5i). We conclude that optogenetic inhibition of climbing fiber signaling blocks learning to a natural, sensory airpuff-US.

### Subtle reductions in climbing fiber signaling eliminate eyeblink conditioning

The simultaneous global silencing of climbing fibers through optogenetic inhibition that we used in Fig. 5 is a dramatic manipulation that could have unexpected consequences for the olivocerebellar circuit. Our final experiment, however, provided additional, unexpected evidence that intact climbing fiber signaling is essential for associative cerebellar learning under natural conditions.

Surprisingly, we found that the CF-ChR2-expressing animals from Fig. 1, who learned well to an optogenetic US, were unable to learn in traditional eyeblink experiments using a sensory airpuff-US, even in the absence of any laser stimulation (CF-ChR2-puff; Fig. 6a-d); %CR at last learning session: CF-ChR2-puff vs airpuff-US controls in Supp. 1o, P=1.98e-06***, Student’s t-Test. In other words, simply expressing ChR2 in climbing fibers completely blocked normal behavioral learning. This surprising result held true despite the facts that, as we have already shown: 1) ChR2-expression was specific to climbing fibers in these mice (Fig. 1c; Supp. Fig. 1). 2) Spontaneous Purkinje cell complex spikes were generally observed in these animals (Fig. 1d-f). 3) Complex spikes were readily evoked by CF-optogenetic stimulation (Fig. 1d-f). 4) The mice displayed intact behavioral unconditioned responses (blinks) to the airpuff (Fig. 6c), indicating intact sensory processing. And of course, 5) CF-ChR2 animals had learned well to an optogenetic CF-ChR2-US (Fig. 1j,k).

**Fig. 6.**
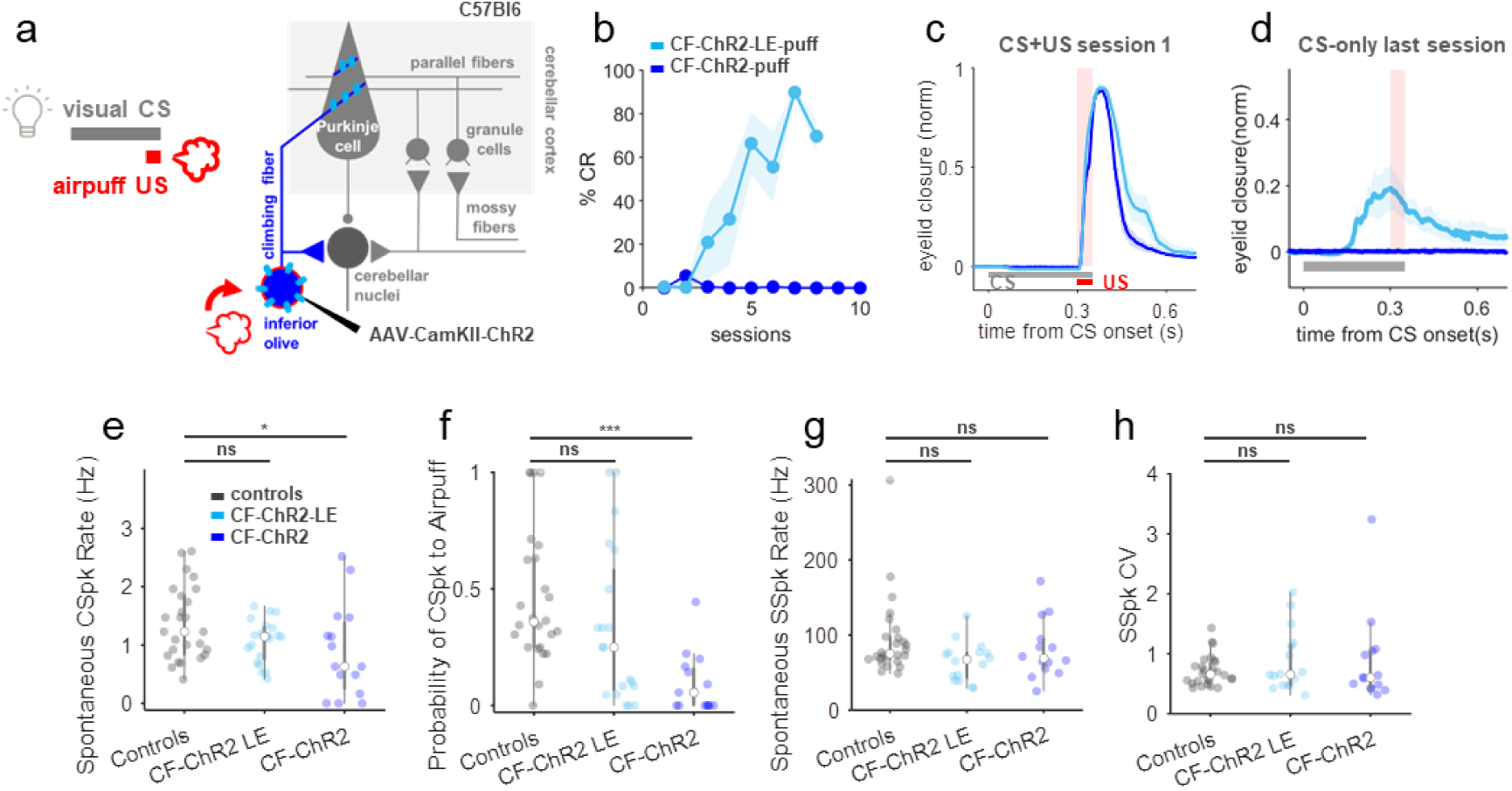
Moderate ChR2 expression is associated with subtle reductions in climbing fiber signaling and abolishes learning to a sensory US. **a**, Experimental scheme. A visual CS was paired with a sensory airpuff-US in a traditional classical conditioning experiment in CF-ChR2 animals. **b**, CF-ChR2-puff animals, without any photostimulation, were unable to learn to an air-puff US (blue, N=6 animals), but lowering ChR2 expression recovered learning (CF-ChR2-LE-puff, light blue, N=4 mice). %CR at last learning session: CF-ChR2-puff vs CF-ChR2-LE-puff, P= 7.75e-05***, Student’s t-Test. **c**, Animals with both expression levels exhibited robust UR blinks on CS+US trials (CF-ChR2-puff, blue, N=6 mice and CF-ChR2-LE-puff, light blue, N=4 mice). **d**, Average eyelid traces from CS-only trials of the last training session reveal no learning in CF-ChR2-puff animals (blue, N=6 mice; CF-ChR2-LE-puff, light blue, N=4 mice). **e**, Spontaneous CSpk firing rate for each Purkinje cell recorded from control (black), CF-ChR2-LE (light blue) and CF-ChR2 (blue) mice; controls vs CF-ChR2, P= 0.04*, Student’s t-Test; controls vs CF-ChR2-LE, P= 0.24n.s., Student’s t-Test. **f**, Probability of an airpuff-evoked complex spike for each Purkinje cell recorded; controls vs CF-ChR2, P= 0.00003***, Student’s t-Test; controls vs CF-ChR2-LE, P= 0.16n.s., Student’s t-Test. **g,h**, SSpk statistics for each Purkinje cell recorded from control (black), CF-ChR2-LE (light blue) and CF-ChR2 (blue) mice. **g**, SSpk spontaneous firing rate; controls vs CF-ChR2, P= 0.41n.s., Student’s t-Test; controls vs CF-ChR2-LE, P= 0.07n.s., Student’s t-Test. **h**, SSpk coefficient of variation; controls vs CF-ChR2, P= 0.33n.s., Student’s t-Test; controls vs CF-ChR2-LE, P= 0.26n.s., Student’s t-Test.

Although we had used standard parameters for viral ChR2 expression^28,48,79^ and there was no obvious anatomical, physiological, or behavioral indication of ChR2 overexpression, we next asked whether lower levels of ChR2-expression in climbing fibers (CF-ChR2-LE; Fig. 1g-l) could restore learning to a sensory US. Remarkably, this 5-fold reduction of viral titer fully restored the ability to learn to a sensory airpuff-US (Fig. 6a-d); %CR at last learning session: CF-ChR2-LE-puff vs airpuff-US controls in Supp. 1o, P=0.12n.s., Student’s t-Test.

To understand how simply expressing ChR2 at moderate levels in climbing fibers could have such a striking and selective impact on learning to a natural sensory US, we quantitatively compared electrophysiological recordings from Purkinje cells in CF-ChR2, CF-ChR2-LE, and control mice (Fig. 6e-h; Supp. Fig. 3). We analyzed spontaneous simple and complex spikes as well as responses to airpuff stimuli delivered to the eye. In control conditions, complex spikes are relatively infrequent, with a low average spontaneous firing rate, and substantial variation across cells (Fig. 6e). We observed subtly lower spontaneous complex spike rates in Purkinje cells of CF-ChR2 mice compared to controls, while no significant reduction was observed in CF-ChR2-LE mice (Fig. 6e). Remarkably, we also found that some (4/15) units with moderate CF-ChR2 expression that showed clear, short-latency complex spikes upon CF-ChR2 stimulation did not exhibit any spontaneous complex spikes throughout the duration of our recordings (Fig. 6e). This surprising finding indicates that the common method of identifying Purkinje cells based on the presence of spontaneous complex spikes would obscure the consequences of CF-ChR2 expression for climbing fiber-Purkinje cell transmission.

We next analyzed the patterns of activity evoked by a sensory airpuff-US (Supp. Fig. 3a-f). There was a dramatic reduction in the probability of complex spikes evoked by an airpuff stimulus in CF-ChR2 mice (Fig. 6f). This was true across the population of Purkinje cells we recorded from these mice, including those with normal spontaneous complex spike rates (Supp. Fig. 3g). In contrast, no systematic reduction in airpuff-evoked complex spiking was observed in the lower expression CF-ChR2-LE animals (Fig. 6f, Supp. Fig. 3h). Further, on trials in which an airpuff did evoke Purkinje cell complex spikes, the responses were delayed in Purkinje cells recorded from CF-ChR2, but not CF-ChR2-LE mice (Supp. Fig. 3 d,f,i).

Importantly, none of the differences in complex spiking observed in CF-ChR2 animals were associated with differences in Purkinje cell simple spike statistics, including average SS firing rate, coefficient of variation, or the pause in simple spikes following a complex spike (Fig. 6g,h, Supp. Fig. 3j). We also found no differences in the number of ‘spikelets’ within each CSpk waveform (Supp. Fig. 3k,l), or in the proportion of CSpks occurring within 200 ms of each other (CSpk ‘doublets’^80^; Supp. Fig. 3m).

## DISCUSSION

The climbing fiber hypothesis for learning has dominated the cerebellar field for over 50 years^1–3^, yet definitive proof – or disproof – has remained elusive. Conflicting evidence, competing models, and insufficiently precise tools for neural circuit dissection have sowed substantial controversy and confusion. In particular, while multiple experimental approaches have yielded data consistent with the theory, others have provided support for an alternative model, in which Purkinje cell simple spike modulation, rather than complex spikes, provides critical instructive signals for learning^4,30,36,43^. Moreover, while sensorimotor errors that drive behavioral learning are often reflected in climbing fiber-driven Purkinje cell CSpk activity, the correlational nature of most of these studies, combined with the unusual spiking statistics of complex spikes, have complicated a definitive interpretation of CSpks as instructive signals^34^. Here we systematically manipulated distinct circuit elements to dissociate climbing fiber-driven complex spike signaling from Purkinje cell simple spike modulation and reflexive movements (Fig. 7). Our findings reveal excitatory climbing fiber inputs as necessary and sufficient instructive signals for associative cerebellar learning.

**Fig. 7.**
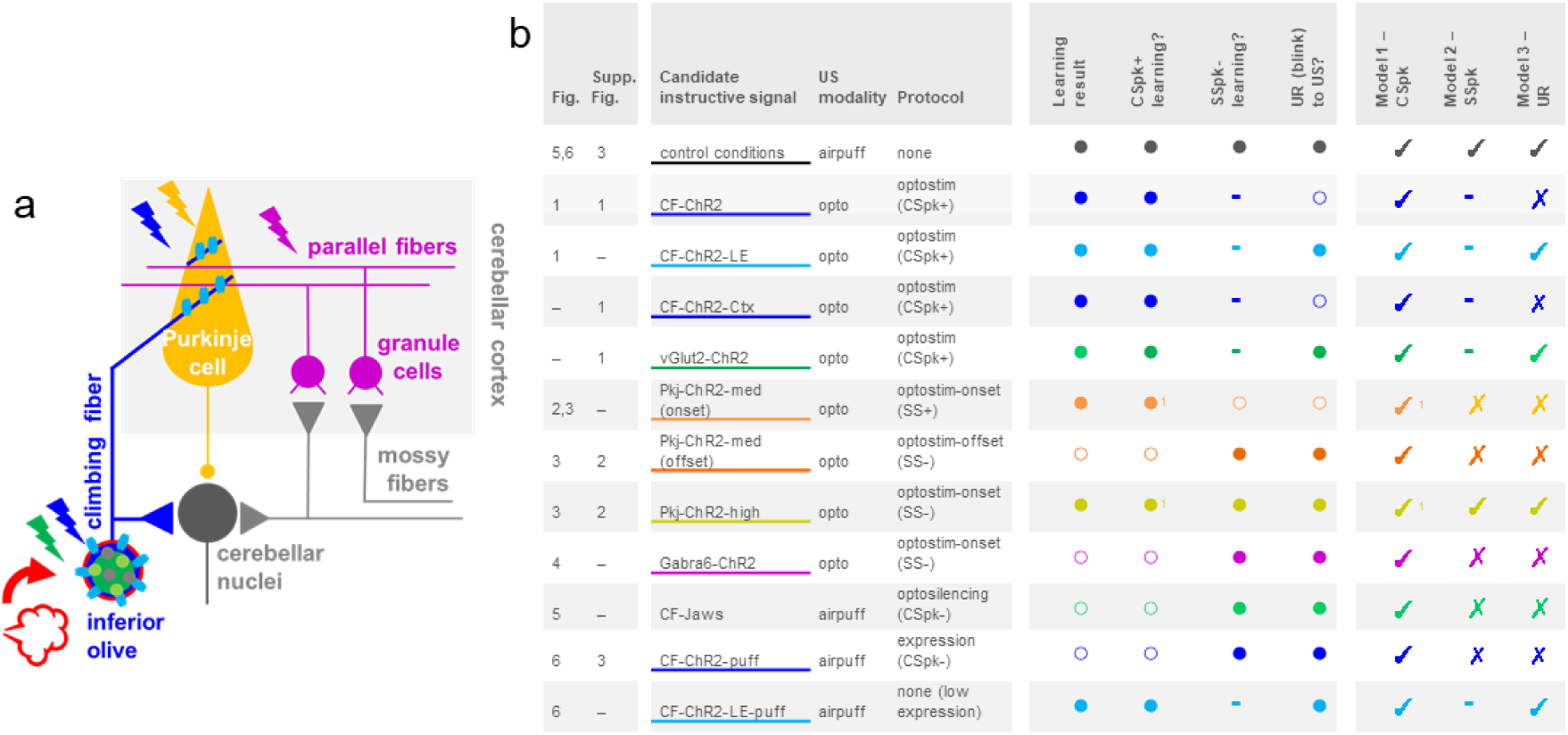
Summary of candidate instructive signals tested in the study and explanatory power of three models for cerebellar learning. **a**, Cerebellar circuit for eyeblink conditioning, highlighting the different strategies used in the study. **b**, Summary table indicating the candidate instructive signal evaluated with each experiment, ordered and color-coded by Figure number. For each candidate tested, the presence/ absence of robust learning is indicated, together with the predictions of blink-, CSpk-, or SSpk-related models for learning. The last 3 columns assess the congruence between each model’s prediction and the learning result that was observed. Closed circles represent ‘yes’, open circles represent ‘no’.

As has been recently shown for VOR adaptation^28,29,48^, we found that optogenetic stimulation of either climbing fibers or Purkinje cells can substitute for a sensory US to drive eyeblink conditioning (Figs. 1-2, Supp. Fig. 1). In both cases, learning was independent of an evoked blink (Figs. 1-3, Supp. Figs. 1-2). However, additional experiments varying opto-Pkj-US laser intensity and duration revealed that learning to a Purkinje cell optogenetic US was temporally coupled to optogenetic US onset, regardless of the direction of Purkinje cell simple spike modulation, or the timing of an evoked blink (Fig. 3). Further experiments in which Purkinje cell simple spike modulation was achieved indirectly, through optogenetic stimulation of granule cells, also failed to induce learning (Fig. 4). These findings suggest that optogenetic stimulation of Purkinje cells likely drives learning through the generation of complex spike-like dendritic calcium signals^48^ (Supp. Fig. 2i), rather than through modulation of simple spike output. This could then also explain why CF-ChR2 expressing animals cannot access Purkinje cell instructive signals to achieve even minimal learning, as we showed in Fig. 6.

Beyond demonstrating their sufficiency as instructive signals for learning, multiple aspects of our data point to the necessity of intact climbing fiber signaling for delay eyeblink conditioning. First, Jaws-mediated optogenetic inhibition of climbing fibers specifically during the presentation of an airpuff-US completely abolished learning (Fig. 5). But perhaps the strongest evidence for the necessity of climbing fiber instructive signals came from our unexpected finding that simply expressing ChR2 in climbing fibers – in the absence of any optical stimulation – reduced complex spike probability and completely obliterated learning to an airpuff-US (Fig. 6). The complete absence of learning to a sensory US in these animals was particularly surprising given the relative subtlety of the effects on complex spiking, and the ability of these mice to learn to an optogenetic CF-US (Fig. 1).

### Mechanistic considerations

The exact mechanism of suppression of climbing fiber signaling by ChR2 expression remains to be determined. Our results point to a decrease in action potential generation within climbing fibers themselves, rather than the postsynaptic generation of complex spikes within Purkinje cells, since the probability of complex spiking was decreased, particularly in response to a sensory US, while complex spike waveforms were not affected (Supp. Fig. 3l) and the ability to induce complex spikes with optogenetic stimulation of climbing fibers was spared. Such changes could also be associated with decreased synchrony across the climbing fiber population. Further, decreases in climbing fiber activity may also alter the likelihood that climbing fibers fire in bursts, or ‘doublets’^80^ under some conditions, although we did not see evidence for this in spontaneous complex spikes (Supp. Fig. 3m), and the drastic reduction in evoked complex spiking that we observed made it impossible to address the potential further contribution of such a mechanism.

One of the most striking aspects of the ChR2 expression effect was its exquisite sensitivity to expression levels – a 5-fold reduction in viral titer was enough to restore both normal complex spiking and learning to an airpuff-US. It is well known that viral gene delivery^81^ and expression of ChR2 and other membrane proteins can alter neuronal morphology and physiology^82–85^ in ways that are still not fully understood. It is possible that IO neurons may be particularly vulnerable, for example due to their high levels of electrical coupling^59,86,87^, which could explain the failure of many previous attempts to target climbing fibers^25^. The use of AAV^81^ and/or of the CaMKIIa promoter may also have contributed, for instance by driving particularly strong expression levels^84^ or through perturbing endogenous CaMKII function in the IO^88^. However, the transgene itself appeared to be critical, as we did not observe a similar phenomenon when using the AAV with the CaMKIIa promoter to drive Jaws expression (Fig. 5h).

The discovery that small changes in ChR2 expression levels can have drastic behavioral consequences has important implications for experiments using optogenetic circuit dissection more broadly. For our purposes, climbing fiber ChR2 expression provided an unexpectedly powerful and selective tool for reducing evoked complex spikes, without affecting Purkinje cell simple spiking and while only subtly reducing spontaneous complex spiking. Still, the effects on complex spiking that we observed were not immediately obvious (see Fig. 1), and depended on comprehensive quantitative analysis that was only possible because cell-type specific activity patterns in the cerebellar circuit and their relationship to relevant sensorimotor signals have previously been exceptionally well-characterized. Thus, while we were able to exploit this unexpected effect as an unparalleled opportunity to assess the contributions of evoked climbing fiber signaling to cerebellar learning, our findings also highlight the major challenge of identifying circuit tools that allow neuroscientists to cleanly isolate and manipulate specific neural signals within complex networks.

### Conclusion and outlook for cerebellar learning

Taken together, our results reconcile many previous, apparently contradictory, findings and suggest that climbing fiber-driven complex spike events provide essential instructive signals for cerebellar learning (Fig. 7).

Our findings also raise important questions about how sensorimotor errors are encoded in the cerebellum to support a full range of cerebellum-dependent behaviors. In particular, it is possible that parallel fiber inputs may provide instructive signals independent of climbing fiber input in some cases. For instance, whole body movements like locomotion generate robust activation of mossy fiber inputs^89,90^. There is evidence that coincident input from spatially clustered parallel fibers can elicit dendritic calcium events in Purkinje cells^91–93^, which could drive cerebellar plasticity in the absence of climbing fiber inputs^94,95^. Although we were not able to induce such an effect via optogenetic stimulation of granule cells (Fig. 4), it remains possible that during some forms of cerebellum-dependent learning, such as motor adaptation^28,96–98^, sufficiently high levels of parallel fiber activation could instruct parallel fiber plasticity.

Similarly, while our results reveal a necessary role for climbing fiber-driven Purkinje cell complex spikes, they do not rule out a possible role for additional plasticity mechanisms in the cerebellar nuclei^99^, which may be important for some forms or components of learning, for instance across time scales ^4,5,100–103^. Previous work has suggested that cerebellar learning may consist of multiple stages, with initial learning in the cerebellar cortex (driven mainly by CF inputs) leading to changes in Purkinje cell output that then sculpt plasticity in the cerebellar nuclei^4,5,100^. The relative contributions of cortical vs. nuclear plasticity may vary across stages of learning, or for different forms of cerebellar learning that progress on different time scales – from short-term motor adaptation over seconds and minutes^17,98,101^, to eyeblink conditioning which takes days^102^, to long-term motor adaptation after prolonged wearing of prism goggles^103^, for example.

Regardless of possible contributions from additional mechanisms, our findings establish an absolute requirement for climbing fiber instructive signals in associative cerebellar learning, and suggest that initial complex spike-driven plasticity could be an essential prerequisite for later stages of cerebellar learning to proceed.

## METHODS

### Animals

All procedures were carried out in accordance with the European Union Directive 86/609/EEC and approved by the Champalimaud Centre for the Unknown Ethics Committee and the Portuguese Direção Geral de Veterinária (Ref. No. 0421/000/000/2015 and 0421/000/000/2020). Mice were kept on a reversed 12-h light/12-h dark cycle with food and water ad libitum. All procedures were performed in male and female mice of approximately 12– 14 weeks of age.

#### Mouse lines

WT C57BL/6J mice were obtained from The Jackson Laboratory (strain #000664). Selective ChR2 expression in Purkinje cells (*L7-Cre;ChR2*; Figs. 2, 3, Supp. Fig 2), granule cells (*Gabra6-Cre;ChR2*; Fig. 4), and glutamatergic neurons within the IO (*vGlut2-Cre;ChR2*; Supp. Fig. 1) were obtained by crossing specific Cre driver lines with ChR2-EYFP-LoxP mice (strain #012569 from The Jackson Laboratory^104^) to generate cell-type specific ChR2-expressing transgenic animals (Supp. Table 1). Cre lines were: For Purkinje cells, L7-Cre strain #004146 from The Jackson Laboratory^53,105^; For granule cells, Gabra6-Cre (MMRRC 000196-UCD^53,54,106,107^); For glutamatergic neurons (within the IO), vGlut2-Cre (strain #016963 from The Jackson Laboratory^60,108,109^).

### Surgical procedures

For all surgeries, animals were anesthetized with isoflurane (4% induction and 0.5 – 1.5% for maintenance), placed in a stereotaxic frame (David Kopf Instruments, Tujunga, CA) and a custom-cut metal head plate was glued to the skull with dental cement (Super Bond – C&B). At the end of the surgery, mice were also administered a non-steroidal anti-inflammatory and painkiller drug (Carprofen). After all surgical procedures, mice were monitored and allowed ∼1-2 days of recovery.

#### Viral injections

Climbing fibers (CF) were targeted^28,48^ by injecting 250nl of AAV1.CaMKIIa.hChR2(H134R)-mCherry.WPRE.hGH (Addgene#26975^110^) or AAV8.CamKII.Jaws-KGC.GFP.ER2-WPRE.SV40 (UPen#AV-8-PV3637^78^) at the left dorsal accessory inferior olive (IO), which has been previously implicated in eyeblink conditioning (RC - 6.3, ML -0.5, DV 5.55^25,26,28^). For the ChR2 virus, we initially diluted the stock virus 1:10 in ACSF to yield a final titer of 1,31×10^12 GC/ml, in line with previous studies^28^. For the low-expression conditions we diluted the virus an additional 5x to yield a final titer of 2,62×10^11 GC/ml. The Jaws virus was diluted in a ratio of 1:10 in ACSF to yield a final titer of 1,47×10^12 GC/ml. CF-ChR2 and CF-Jaws mice started the behavioral and electrophysiological experiments 6 weeks after injection to allow time for virus expression and stabilization^28^.

**Supp. Table 1.** Summary of genetic and anatomical targeting approaches used for selective targeting of individual cerebellar circuit elements.

| Fig # | Experimental condition | Expression strategy | Mouse genotype | Virus | Promoter | Cell type targeted | Fiber placement |
| --- | --- | --- | --- | --- | --- | --- | --- |
| 1, 6, S1, S3 | **CF-ChR2** | viral | wt(C57Bl6) | AAV-CaMKII-ChR2 ($10^{12}$) | CaMKII | **CF** | **Ctx / IO** |
| 1, 6, S1, S3 | **CF-ChR2-LE** | viral -Low Expression | wt(C57Bl6) | AAV-CaMKII-ChR2 ($10^{11}$) | CaMKII | **CF** | **IO** |
| S1 | **vGlut2-ChR2-IO** | transgenic | vGlut2-Cre;ChR2 |  | **vGlut2** | IO-glut | **IO** |
| 2, 3, S2 | **Pkj-ChR2** | transgenic | L7-Cre;ChR2 |  | **L7** | **Pkj** | **Ctx** |
| 4 | **Gabra6-ChR2-Ctx** | transgenic | Gabra6-Cre;ChR2 |  | **Gabra6** | granule cells | **Ctx** |
| 5 | **CF-Jaws** | viral | wt(C57Bl6) | AAV-CaMKII-Jaws ($10^{12}$) | CaMKII | **CF** | **IO** |

For optogenetic manipulations (Supp. Table 1), optical fibers with 100μm core diameter, 0.22 NA (Doric lenses, Quebec, Canada) were lowered into the brain through small craniotomies performed with a dental drill, and positioned at either the right cerebellar cortical eyelid region (RC -5.7, ML +1.9, DV -1.5)^56,58,59^ or at the left dorsal accessory inferior olive (IO; RC -6.3, ML - 0.5, DV -5.5), which has been previously implicated in eyeblink conditioning^25,26,111^. Correct fiber placement in both the cerebellar cortex and IO was functionally verified before experiments by the presence of an evoked eyeblink in the right eye in response to moderate intensity laser stimulation (when possible; see below) and subsequently confirmed histologically. Only animals with good opsin expression and precise fiber targeting were kept in the study.

For in vivo electrophysiological recordings, a disposable 3 mm biopsy punch was used to perform a craniotomy over the right cerebellar cortical eyelid region (RC -5.7, ML +1.9^56,58,59^. The craniotomy was covered with a 3mm glass coverslip with 4 small holes where the electrode could pass through, and then by silicon based elastomer (Kwik-cast, WPI) that was easily removed just before recording sessions.

### Behavioral procedures

The experimental setup for eyeblink conditioning was based on previous work^53,54^. For all behavioral experiments, mice were head-fixed and walking on a Fast-Trac Activity Wheel (Bio-Serv). A DC motor with an encoder (Maxon) was used to externally control the speed of the treadmill. Mice were habituated to the behavioral setup for at least 4 days prior to training, until they walked normally at the target speed of 0.1m/s and displayed no external signs of distress. Eyelid movements of the right eye were recorded using a high-speed monochromatic camera (Genie HM640, Dalsa) to monitor a 172×160 pixel region at 900fps. We visually monitored whole-body movements via webcam continuously throughout each experiment. Custom-written LabVIEW software, together with a NI PCIE-8235 frame grabber and a NI-DAQmx board (National Instruments), was used to synchronously trigger and control the hardware.

Acquisition sessions consisted of the presentation of 90 CS+US paired trials and 10 CS-only trials. The 100 trials were separated by a randomized inter trial interval (ITI) of 10-15s. Unless otherwise stated, CS and US onsets on CS+US paired trials were separated by a fixed inter-stimulus interval (ISI) of 300ms and both stimuli co-terminated. The CS was a white light LED positioned ∼3cm directly in front of the mouse. The sensory unconditioned stimulus (US) was an airpuff (40psi, 50ms) controlled by a Picospritzer (Parker) and delivered via a 27G needle positioned ∼0.5cm away from the cornea of the right eye of the mouse. Airpuff direction was adjusted for each session of each mouse so that the US elicited a strong reflexive eye blink unconditioned response (UR).

#### Behavioral analysis

Videos from each trial were analyzed offline with custom-written MATLAB (MathWorks) software^53^. Distance between eyelids was calculated frame by frame by thresholding the grayscale image and extracting the minor axis of the ellipse that delineated the eye. Eyelid traces were normalized for each session, from 0 (maximal opening of the eye throughout the session) to 1 (full eye closure achieved under airpuff treatment). Trials were classified as containing CRs if an eyelid closure with normalized amplitude >0.1 occurred >100ms after CS onset and before US onset.

### Optogenetic stimulation and inhibition

Light from 473 or 594 nm lasers (LRS-0473or LRS-0594 DPSS, LaserGlow Technologies; excitation and inhibition, respectively) was controlled with custom-written LabView code. Predicted irradiance levels for the 100um diameter, 0.22NA optical cannulas used in our study were calculated using the online platform: https://web.stanford.edu/group/dlab/optogenetics. All laser powers are comparable to those of previous studies^28,29,47,48,53,56,112^.

For Pkj-ChR2 (Figs. 2, 3 and Supp. Fig. 2), Gabra6-ChR2 (Fig. 4) and vGlut2-ChR2-IO (Supp. Fig. 1) experiments, laser power was adjusted for each mouse and controlled for each experiment using a light power meter (Thorlabs) at the beginning and end of each session. For the Pkj-ChR2 experiments of Fig. 2, Fig. 3a-i, and Supp. Fig 2b-d, laser intensity was adjusted to elicit an intermediate eyelid closure, and no other body movements, at stimulus offset (1-3mW, max irradiance of 95.5mW/mm^2. For the Pkj-ChR2-high (blink at laser onset) experiments of Fig. 3j-m and Supp. Fig. 2e-h powers ranged from 8-12mW, max irradiance of 381.8mW/mm^2. For the Gabra6-ChR2 experiments of Fig. 4, intensities were up to 6mW, irradiance of 190.9mW/mm^2 (causing a blink at laser onset and no other body movements). For the vGlut2-ChR2-IO of Supp. Fig. 1, laser power was adjusted to elicit a blink (and no other body movements) at laser onset (vGlut2-ChR2-IO: 1-3.3mW, max predicted irradiance of 105mW/mm^2).

For the CF-ChR2 experiments of Fig. 1, since these animals did not blink (or present any other body movements) to laser stimulation (Fig. 1i), the power was set to 6mW (max predicted irradiance of 190.9mW/mm^2). This power was confirmed with electrophysiology to reliably drive Purkinje cell complex spikes. The same power was also used for all CF-ChR2-LE experiments; which exhibited a small eyelid twitch (and no other body movements) in response to laser stimulation (Fig. 1i).

When optogenetic stimulation was substituting for a sensory US, where possible we adjusted the timing (onset and duration) of the laser stimulation so that the reflexive blinks would most closely match those elicited by a sensory US (50ms airpuff delivered to the eye). For Pkj-ChR2 and Gabra6-ChR2 experiments, 100ms laser stimulation best elicited a blink similar to that of the airpuff. Because Pkj-ChR2-med stimulation elicits a blink at the offset of laser stimulation, whereas GC-ChR2 elicits a blink at the onset of laser stimulation (due to Purkinje cell inhibition via molecular layer interneurons), the onset of the laser stimulation was also adjusted specifically for those experiments. For the CF-ChR2 experiments of Fig. 1, since there was no laser-driven blink (Fig. 1i), we kept the 100ms laser duration and matched the timing of laser stimulation/ complex spike onset.

For the CF-Jaws experiments of Fig. 5, since these animals did not blink or present any other body movement to laser inhibition, the power was set to 6mW (max predicted irradiance of 190.9mW/mm^2). This power was confirmed with electrophysiology to reliably block airpuff-driven Purkinje cell complex spikes. For these inhibition experiments, laser started at time of airpuff onset and laser duration was randomized between 300-400ms in order to avoid consistently timed rebound excitation^78^.

### Electrophysiological recordings

All recordings were performed in vivo, in awake mice. Cell-attached single-cell recordings were made using long-shanked borosilicate glass pipettes (Warner Instruments) pulled on a vertical puller (Narishige PC-100) and filled with saline solution (0.9% NaCl, typical resistances between 4-5 MΩ). An Optopatcher (A-M Systems) was used for simultaneous optogenetic stimulation and electrophysiological recordings. Laser light (with the same blue laser used for the behavioral optogenetic manipulations) was transmitted through an optic fiber (50 μm core diameter) inserted inside the glass pipette until it could fit, ∼5 mm from the tip. The Optopatcher was oriented towards the cerebellar eyeblink region with a motorized 4-axis micromanipulator (PatchStar, Scientifica). Craniotomies were filled with saline and connected to the ground reference using a silver-chloride pellet (Molecular Devices).

Recordings were performed with a Multiclamp 700B amplifier (Axon Instruments) in its voltage-clamp configuration, with a gain of 0.5 V/nA and low-pass Bessel filter with 10 kHz cut-off. The current offset between the interior and exterior of the pipette was always kept neutral to avoid passive stimulation of the cells. All recordings were sampled at 25kHz from the output signal of the amplifier using a NI-DAQmx board and Labview custom-software. Purkinje cells were generally identified by the presence of complex spikes. In cells that exhibited spontaneous complex spikes (a typical criterion for identifying Purkinje cells in electrophysiological recordings), we observed subtly lower spontaneous complex spike rates in Purkinje cells of CF-ChR2 mice compared to controls. Because of this, for all CF-ChR2 (and CF-ChR2-LE) experiments, Purkinje cells were identified based on the presence of a laser-triggered complex spike rather than spontaneous complex spikes, to avoid selection bias resulting from the absence of spontaneous complex spiking (Fig. 6). Spikes were sorted offline using custom Python code for simple spikes and a modified Un’Eye neural network^113^ for complex spikes.

### Histology

All experiments included histological verification of injection and fiber placement and transgene expression levels. After the experiments, animals were perfused transcardially with 4% paraformaldehyde and their brains removed. Brain sections (50um thick) were cut in a vibratome and stained for Purkinje cells (with chicken anti-calbindin primary antibody #214006 SYSY, and anti-chicken Alexa488 #703-545-155 or Alexa594 #703-545-155 secondary antibodies from Jackson Immunoresearch) and for cell viability (with DAPI marker). Brain sections were mounted on glass slides with mowiol mounting medium, and imaged with 5x, 10x or 20x objectives. Brain slices from experiments where Climbing Fibers were targeted were also imaged with an upright confocal laser point-scanning microscope (Zeiss LSM 710), using a 10x or 40x objective.

### Statistical analysis

Data are reported as mean ± s.e.m., and statistical analyses were performed using the Statistics toolbox in MATLAB. Two-sample two-tailed or paired Student’s t-Tests (specified in each case) were performed for all comparisons unless otherwise indicated. Differences were considered significant at *P<0.05, **P<0.01, and ***P<0.001. No statistical methods were used to predetermine sample sizes; sample sizes are similar to those reported in previous publications^53,54,56^. Data collection and analysis were not performed blind to the conditions of the experiments. Mice were randomly assigned to specific experimental groups without bias.

## ACKNOWLEDGEMENTS

We thank T. Pritchett and A. Machado for maintenance of mouse lines and C. Almeida for technical assistance with some experiments. We thank Champalimaud Research Vivarium Staff, Histology, Microscopy, Hardware and Sci-Comm Platforms for technical support and A. Gonçalves for assistance with microscopy. We are grateful to the Carey lab and other members of the Champalimaud Neuroscience Program for helpful discussions throughout the project and to H. Marques, C. Hérent, and C. Albergaria for critical feedback on the manuscript. This work was supported by fellowships from the Portuguese Fundação para a Ciência e a Tecnologia (FCT) #BD/105949/2014 (to NTS) and #BPD109659/2015 (to DLP), Bial Foundation Bursary #74/14 (to DLP), and grants to MRC from the Howard Hughes Medical Institute #55007413, FCT #PTDC/MED_NEU/30890/2017, and European Research Council #866237. Additional support was provided by Congento LISBOA-01–0145-FEDER-022170, co-financed by FCT (Portugal) and Lisboa2020 under PORTUGAL2020 agreement.

## SUPPLEMENTAL FIGURES

**Supp. Fig. 1.**
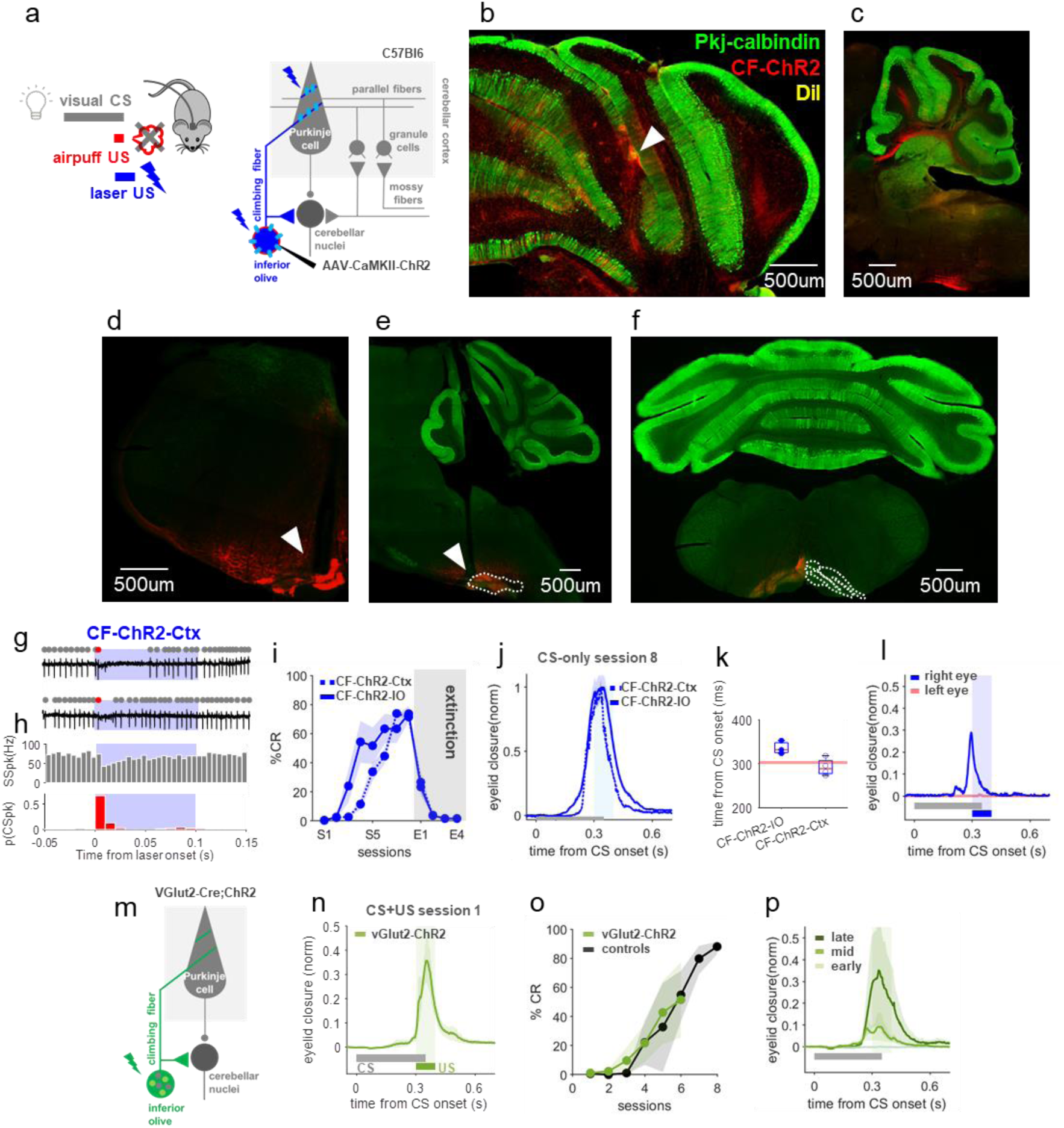
Multiple genetic and anatomical strategies for targeting IO neurons. **a**, Experimental scheme. **b**, Example coronal section showing fiber placement (white arrow), at the eyelid area of the right cerebellar cortex of CF-ChR2 animals. **c**, Example sagittal section showing ChR2 expression in climbing fiber projections to the cerebellar cortex. **d**, Example coronal section showing CF-ChR2 expression and optical fiber placement (white arrow) at the left IO. **e**, Example sagittal section showing CF-ChR2 expression and fiber placement (white arrow) at the left IO. **f**, Example coronal section showing ChR2 expression at the left inferior olive of CF-ChR2 expressing animals. **g**, Two example electrophysiological traces from a Purkinje cell with identified SSpks (grey dots) and CSpks (red) in response to CF-ChR2 laser stimulation in the cerebellar cortex, targeting CF terminals (CF-ChR2-Ctx, blue). **h**, Population histogram of SSpks (grey) and CSpks (red) (n=211 trials, N=15 units from 5 animals). CSpks: spont. vs laser, P=1.82e-06***, Student’s paired t-Test; SSpks: spont. vs laser, **P=0.002, Student’s paired t-Test. **i**, %CR over the course of multiple training (S1-S8) and extinction sessions (E1-E4, where only the visual CS is presented without a US) of animals with CF-ChR2 laser stimulation at climbing fiber somas in the IO (CF-ChR2-IO, solid line, N=3 mice) or terminals at the cerebellar cortex as US (CF-ChR2-Ctx, dotted line, N=4 mice). %CR at last learning session: CF-ChR2-IO vs CF-ChR2-Ctx, P=0.71n.s., Student’s t-Test. **j**, Normalized traces illustrate broader CR timing for the CF-ChR2-IO condition (for the experiments shown in *i*), which corresponds with the more sustained increase in CSpks during optogenetic stimulation in the IO vs ctx (Fig. 1f,h vs Supp. Fig 1h). **k**, Timing of peak eyelid closures (for the last learning session of experiments shown in *i*) are subtly later for the CF-ChR2-IO condition, across mice. Peak eyelid closure time: CF-ChR2-IO vs CF-ChR2-Ctx, P=0.37n.s, Student’s t-Test. Each dot is one mouse and box plots indicate 25th-75th percentiles with whiskers extending to most extreme data point. **l**, Average eyeblink closure traces after eyeblink learning with CF-ChR2 laser stimulation at the cerebellar cortex as an US. Conditioning was unilateral (right conditioned eye in blue, left eye in red; N=2 mice). **m**, Experimental scheme for experiments stimulating glutamatergic IO neurons as a US (vGlut2-ChR2-IO). **n,** Average eyelid closures on CS+US trials in the first training session showing the blink evoked by vGlut2-ChR2-IO laser stimulation (N=3 mice). **o**, %CR across sessions for learning to vGlut2-ChR2-IO laser stimulation (green, N=3 mice) and controls (expressing ChR2 but without laser stimulation, learning to an airpuff-US); %CR at last learning session: vGlut2-ChR2-IO vs controls, P=0.94n.s., Student’s t-Test. **p**, Average eyelid closure traces from CS-only trials of S2, S4 and S6 training sessions for a vGlut2-ChR2-IO-US (N=3 mice).

**Supp. Fig. 2.**
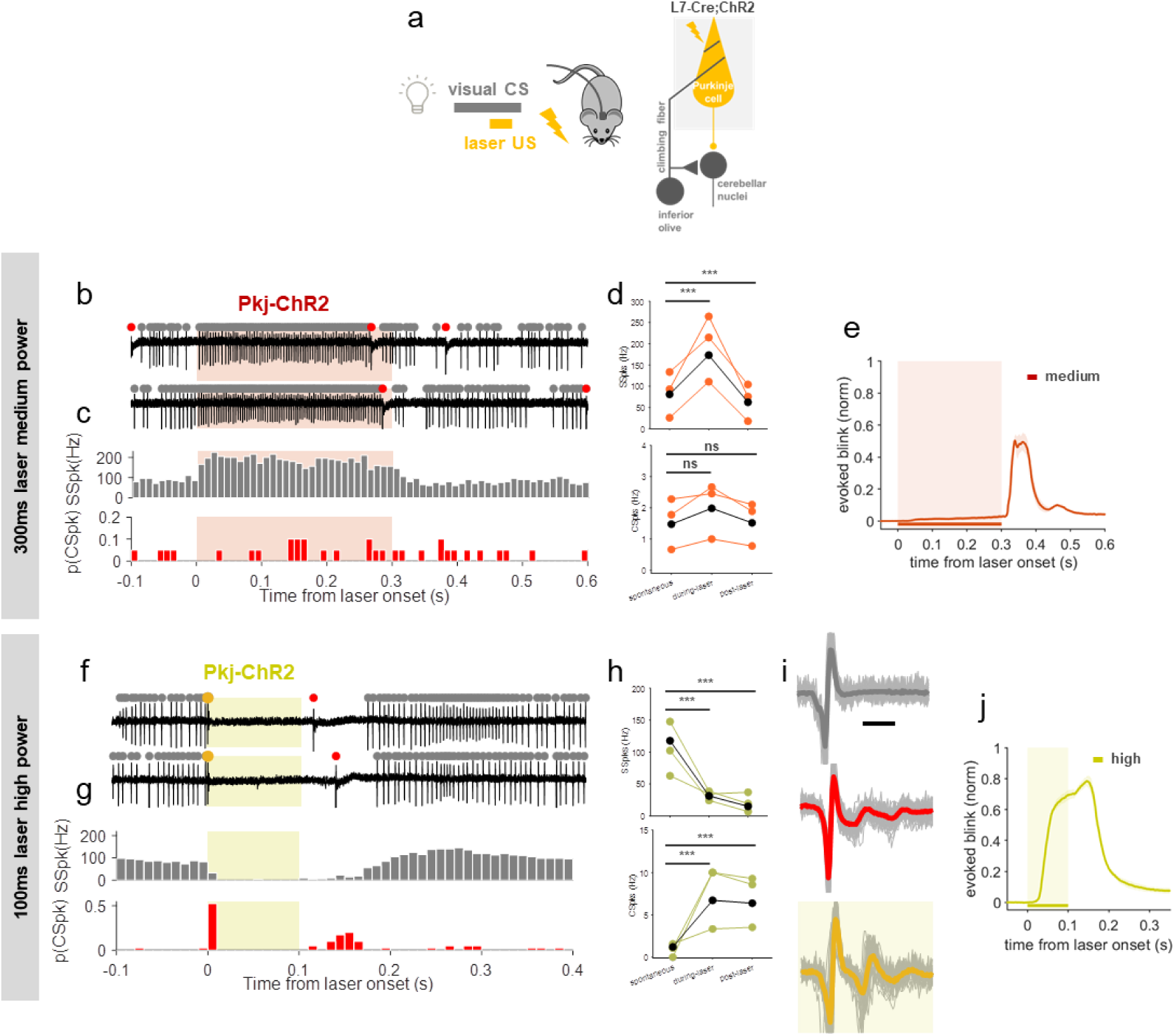
Varying intensity and duration of Purkinje cell optogenetic stimulation to dissociate stimulation onset, simple spike modulation, and evoked blinks. **a**, Experimental scheme for pairing a visual CS with optogenetic Purkinje cell-US. **b,c,** Example electrophysiological traces (b) and histogram (c) from a Purkinje cell with identified SSpk (grey dots) and CSpk (red) in response to 300ms medium intensity Pkj-ChR2 laser stimulation in the cerebellar cortex (Pkj-ChR2-Ctx-med; corresponds to Fig. 3f-i). **d**, SSpk rate (top) and CSpk rate (bottom) pre-, during- and post-laser epochs for the recordings in (b,c) and Fig. 2d (N=3 units from 2 mice). SSpks: spont. vs during laser, P=0. 4.8e-7***, Student’s paired t-Test; spont. vs after laser, P= 0.0003***, Student’s paired t-Test. CSpks: spont. vs during laser, P= 0.22n.s., Student’s paired t-Test; spont. vs after laser, P= 0.87n.s., Student’s paired t-Test. Each orange circle and line represents a unit, linked through conditions; the black solid circles and line represent the average. **e**, Average eyelid closures to 300ms Pkj-ChR2-Ctx medium intensity laser stimulation (N=2 mice, shading represents laser stimulation). **f-j** Higher intensity Pkj-ChR2-Ctx laser stimulation was used to evoke a pause in simple spikes and a short-latency evoked blink at stimulus *onset* (corresponds to Fig. 3j-m). This stimulation elicited electrophysiological signatures of complex spike-like events at laser onset^48^ and, with a longer and more variable delay, at laser offset (likely due to rebound from release of Purkinje cell inhibition via olivo-cerebellar loop^114^. **f,g,** SSpk and CSpk traces and histograms (N=2 units from 2 mice). **h**, Same as in *d*, but for high intensity Pkj-ChR2-Ctx laser stimulation. SSpks: spont. vs during laser, P= 9.9e-17***, Student’s paired t-Test; spont. vs after laser, P= 9.86e-20***, Student’s paired t-Test. CSpks: spont. vs during laser, P= 1.335e-15***, Student’s paired t-Test; spont. vs after laser, P= 2.45e-14***, Student’s paired t-Test. **i,** SSpk (grey) and spontaneous CSpk (red) waveforms. Yellow trace represents complex spike-like events at laser stimulation onset; note the correspondence to spontaneous CSpk waveforms (red). **j**, Pkj-ChR2 laser stimulation at higher intensities yields a blink at stimulus *onset* (N=3 mice, shading represents laser stimulation).

**Supp. Fig. 3.**
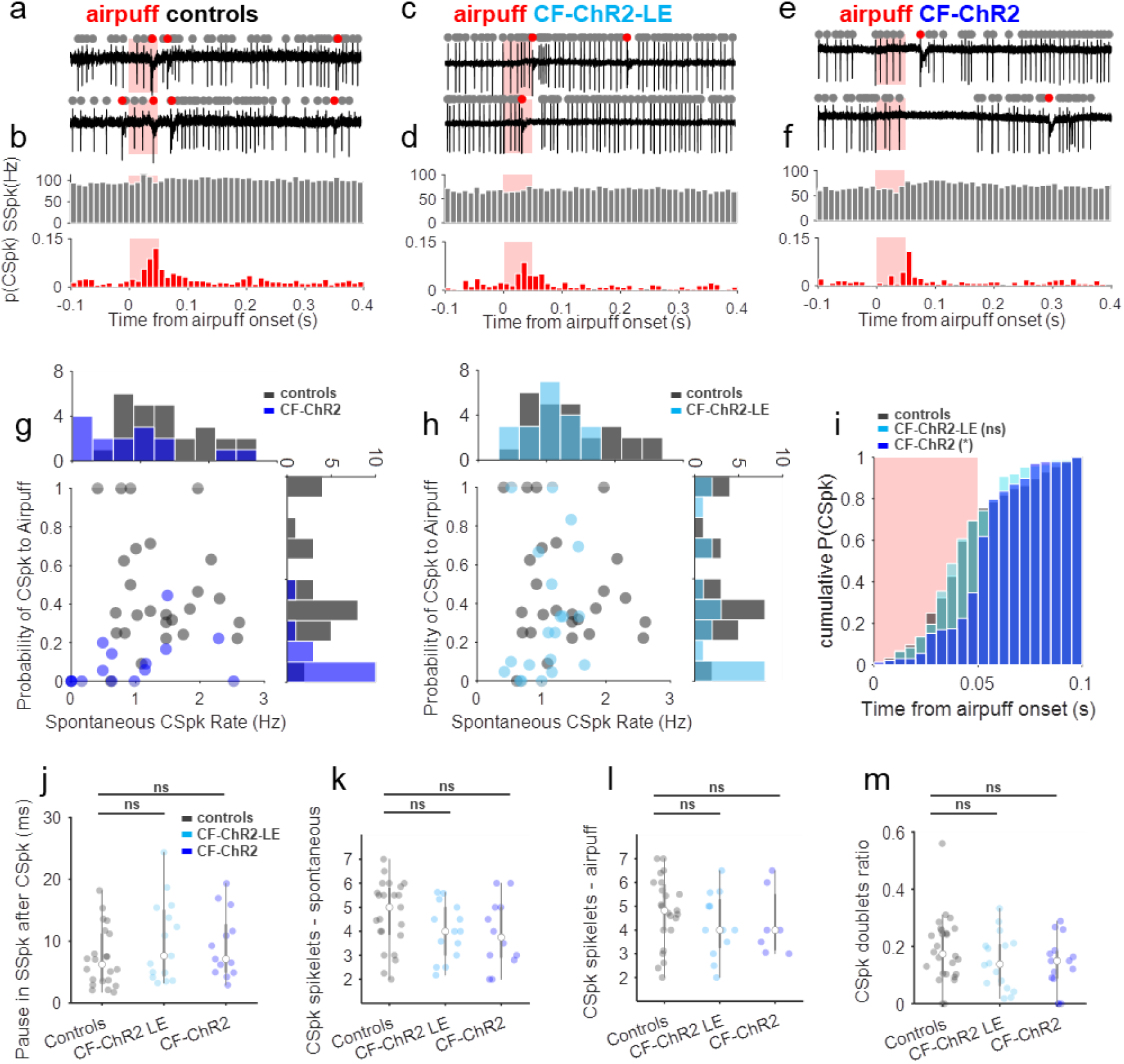
Purkinje cell responses to an airpuff stimulus in controls and CF-ChR2 animals with different expression levels. **a,b**, Example electrophysiological traces and population histograms (N=26 units from 4 mice) of Purkinje cell SSpks (grey dots) and CSpks (red) from control mice in response to an airpuff to the eye. **c,d**, Same as *a,b* but for mice expressing low levels of ChR2 in CFs (CF-ChR2-LE; N=20 units from 4 mice). **e,f**, Same as *c,d* but for CF-ChR2 at standard expression levels (N=15 units from 5 mice). **g**, p(CSpk) to airpuff as a function of spontaneous CSpk rate for each Purkinje cell of controls vs. mice with standard levels of CF-ChR2 expression (CF-ChR2). **h**, Same as *g*, but comparing controls vs. mice with low levels of CF-ChR2 expression (CF-ChR2-LE). **i**, Normalized cumulative histogram of the timing of the first CSpk after airpuff onset (grey: controls N=26 cells from 4 animals; light blue, CF-ChR2-LE, N=20 cells from 4 animals; blue: CF-ChR2 N=15 cells from 5 mice); controls vs CF-ChR2, P= 0.03*, KS-test; controls vs CF-ChR2-LE, P= 0.67n.s., KS-test. Shaded rectangle indicates time of airpuff (red). **j**, Average pause in SSpk activity after a CSpk: controls vs CF-ChR2, P= 0.15n.s., Student’s t-Test; controls vs CF-ChR2-LE, P= 0.24n.s., Student’s t-Test. **k,l**, Average number of spontaneous and airpuff-driven CSpk spikelets, respectively. Spont.: controls vs CF-ChR2, P= 0.1n.s., Student’s t-Test; controls vs CF-ChR2-LE, P= 0.1n.s., Student’s t-Test. Airpuff: controls vs CF-ChR2, P= 0.5n.s., Student’s t-Test; controls vs CF-ChR2-LE, P= 0.3n.s., Student’s t-Test. **m**, Average CSpk doublets (2 CSpks occurring within 200 ms of each other) ratio to total number of CSpks during spontaneous and airpuff epochs; controls vs CF-ChR2, P= 0.14n.s., Student’s t-Test; controls vs CF-ChR2-LE, P= 0.24n.s., Student’s t-Test.

## REFERENCES

1. Marr, D. (1969). A theory of cerebellar cortex. J. Physiol. 202, 437–470. 10.1113/jphysiol.1969.sp008820.

2. Albus, J.S. (1971). A theory of cerebellar function. Math. Biosci. 10, 25–61. 10.1016/0025-5564(71)90051-4.

3. Ito, M. (1972). Neural design of the cerebellar motor control system. Brain Res. 40, 81–84. 10.1016/0006-8993(72)90110-2.

4. Raymond, J.L., Lisberger, S.G., and Mauk, M.D. (1996). The cerebellum: a neuronal learning machine? Science 272, 1126–1131. 10.1126/science.272.5265.1126.

5. Mauk, M.D., and Donegan, N.H. (1997). A model of Pavlovian eyelid conditioning based on the synaptic organization of the cerebellum. Learn. Mem. 4, 130–158. 10.1101/lm.4.1.130.

6. Medina, J.F., Garcia, K.S., Nores, W.L., Taylor, N.M., and Mauk, M.D. (2000). Timing Mechanisms in the Cerebellum: Testing Predictions of a Large-Scale Computer Simulation. J. Neurosci. 20, 5516 LP – 5525. 10.1523/JNEUROSCI.20-14-05516.2000.

7. De Zeeuw, C.I., and Yeo, C.H. (2005). Time and tide in cerebellar memory formation. Curr. Opin. Neurobiol. 15, 667–674. 10.1016/j.conb.2005.10.008.

8. Raymond, J.L., and Medina, J.F. (2018). Computational Principles of Supervised Learning in the Cerebellum. Annu. Rev. Neurosci. 41, 233–253. 10.1146/annurev-neuro-080317-061948.

9. Lisberger, S.G. (2021). The Rules of Cerebellar Learning: Around the Ito Hypothesis. Neuroscience 462, 175–190. 10.1016/j.neuroscience.2020.08.026.

10. Ito, M., and Kano, M. (1982). Long-lasting depression of parallel fiber-Purkinje cell transmission induced by conjunctive stimulation of parallel fibers and climbing fibers in the cerebellar cortex. Neurosci. Lett. 33, 253–258. 10.1016/0304-3940(82)90380-9.

11. Schmolesky, M., Weber, J., de Zeeuw, C., and Hansel, C. (2002). The making of a complex spike: Ionic composition and plasticity. 10.1111/j.1749-6632.2002.tb07581.x.

12. Coesmans, M., Weber, J.T., De Zeeuw, C.I., and Hansel, C. (2004). Bidirectional parallel fiber plasticity in the cerebellum under climbing fiber control. Neuron 44, 691–700. 10.1016/j.neuron.2004.10.031.

13. Carey, M.R. (2011). Synaptic mechanisms of sensorimotor learning in the cerebellum. Curr. Opin. Neurobiol. 21, 609–615. 10.1016/j.conb.2011.06.011.

14. Ramirez, J.E., and Stell, B.M. (2016). Calcium Imaging Reveals Coordinated Simple Spike Pauses in Populations of Cerebellar Purkinje Cells. Cell Rep. 17, 3125–3132. 10.1016/j.celrep.2016.11.075.

15. Gilbert, P.F., and Thach, W.T. (1977). Purkinje cell activity during motor learning. Brain Res. 128, 309–328. 10.1016/0006-8993(77)90997-0.

16. Sears, L.L., and Steinmetz, J.E. (1990). Acquisition of classically conditioned-related activity in the hippocampus is affected by lesions of the cerebellar interpositus nucleus. Behav. Neurosci. 104, 681–692. 10.1037/0735-7044.104.5.681.

17. Medina, J.F., and Lisberger, S.G. (2008). Links from complex spikes to local plasticity and motor learning in the cerebellum of awake-behaving monkeys. Nat. Neurosci. 11, 1185–1192. 10.1038/nn.2197.

18. Rasmussen, A., Jirenhed, D.-A., Wetmore, D.Z., and Hesslow, G. (2014). Changes in complex spike activity during classical conditioning. Front. Neural Circuits 8, 90. 10.3389/fncir.2014.00090.

19. Kimpo, R.R., Rinaldi, J.M., Kim, C.K., Payne, H.L., and Raymond, J.L. (2014). Gating of neural error signals during motor learning. Elife 3, e02076–e02076. 10.7554/eLife.02076.

20. Ohmae, S., and Medina, J.F. (2015). Climbing fibers encode a temporal-difference prediction error during cerebellar learning in mice. Nat. Neurosci. 18, 1798–1803. 10.1038/nn.4167.

21. Stone, L.S., and Lisberger, S.G. (1990). Visual responses of Purkinje cells in the cerebellar flocculus during smooth-pursuit eye movements in monkeys. II. Complex spikes. J. Neurophysiol. 63, 1262–1275. 10.1152/jn.1990.63.5.1262.

22. Kahlon, M., and Lisberger, S.G. (1996). Coordinate system for learning in the smooth pursuit eye movements of monkeys. J. Neurosci. 16, 7270–7283. 10.1523/JNEUROSCI.16-22-07270.1996.

23. Kim, J.J., Krupa, D.J., and Thompson, R.F. (1998). Inhibitory cerebello-olivary projections and blocking effect in classical conditioning. Science 279, 570–573. 10.1126/science.279.5350.570.

24. Medina, J.F., Nores, W.L., and Mauk, M.D. (2002). Inhibition of climbing fibres is a signal for the extinction of conditioned eyelid responses. Nature 416, 330–333. 10.1038/416330a.

25. Kim, O.A., Ohmae, S., and Medina, J.F. (2020). A cerebello-olivary signal for negative prediction error is sufficient to cause extinction of associative motor learning. Nat. Neurosci. 23, 1550–1554. 10.1038/s41593-020-00732-1.

26. Mauk, M.D., Steinmetz, J.E., and Thompson, R.F. (1986). Classical conditioning using stimulation of the inferior olive as the unconditioned stimulus. Proc. Natl. Acad. Sci. U. S. A. 83, 5349–5353. 10.1073/pnas.83.14.5349.

27. Steinmetz, J.E., Lavond, D.G., and Thompson, R.F. (1989). Classical conditioning in rabbits using pontine nucleus stimulation as a conditioned stimulus and inferior olive stimulation as an unconditioned stimulus. Synapse 3, 225–233. 10.1002/syn.890030308.

28. Nguyen-Vu, T.D.B., Kimpo, R.R., Rinaldi, J.M., Kohli, A., Zeng, H., Deisseroth, K., and Raymond, J.L. (2013). Cerebellar Purkinje cell activity drives motor learning. Nat. Neurosci. 16, 1734–1736. 10.1038/nn.3576.

29. Rowan, M.J.M., Bonnan, A., Zhang, K., Amat, S.B., Kikuchi, C., Taniguchi, H., Augustine, G.J., and Christie, J.M. (2018). Graded Control of Climbing-Fiber-Mediated Plasticity and Learning by Inhibition in the Cerebellum. Neuron 99, 999–1015.e6. 10.1016/j.neuron.2018.07.024.

30. Ke, M.C., Guo, C.C., and Raymond, J.L. (2009). Elimination of climbing fiber instructive signals during motor learning. Nat. Neurosci. 12, 1171–1179. 10.1038/nn.2366.

31. Zbarska, S., Bloedel, J.R., and Bracha, V. (2008). Cerebellar Dysfunction Explains the Extinction-Like Abolition of Conditioned Eyeblinks After NBQX Injections in the Inferior Olive. J. Neurosci. 28, 10 LP – 20. 10.1523/JNEUROSCI.3403-07.2008.

32. Schonewille, M., Gao, Z., Boele, H.-J., Vinueza Veloz, M.F., Amerika, W.E., Šimek, A.A.M., De Jeu, M.T., Steinberg, J.P., Takamiya, K., Hoebeek, F.E., et al. (2011). Reevaluating the Role of LTD in Cerebellar Motor Learning. Neuron 70, 43–50. 10.1016/j.neuron.2011.02.044.

33. Popa, L.S., Streng, M.L., Hewitt, A.L., and Ebner, T.J. (2016). The Errors of Our Ways: Understanding Error Representations in Cerebellar-Dependent Motor Learning. The Cerebellum 15, 93–103. 10.1007/s12311-015-0685-5.

34. Streng, M.L., Popa, L.S., and Ebner, T.J. (2018). Complex Spike Wars: a New Hope. The Cerebellum 17, 735–746. 10.1007/s12311-018-0960-3.

35. Sanger, T.D., and Kawato, M. (2020). A Cerebellar Computational Mechanism for Delay Conditioning at Precise Time Intervals. Neural Comput. 32, 2069–2084. 10.1162/neco_a_01318.

36. Miles, F.A., and Lisberger, S.G. (1981). Plasticity in the vestibulo-ocular reflex: a new hypothesis. Annu. Rev. Neurosci. 4, 273–299. 10.1146/annurev.ne.04.030181.001421.

37. Popa, L.S., Hewitt, A.L., and Ebner, T.J. (2012). Predictive and Feedback Performance Errors Are Signaled in the Simple Spike Discharge of Individual Purkinje Cells. J. Neurosci. 32, 15345 LP – 15358. 10.1523/JNEUROSCI.2151-12.2012.

38. Lisberger, S.G., and Fuchs, A.F. (1978). Role of primate flocculus during rapid behavioral modification of vestibuloocular reflex. I. Purkinje cell activity during visually guided horizontal smooth-pursuit eye movements and passive head rotation. J. Neurophysiol. 41, 733–763. 10.1152/jn.1978.41.3.733.

39. Kawato, M., Furukawa, K., and Suzuki, R. (1987). A hierarchical neural-network model for control and learning of voluntary movement. Biol. Cybern. 57, 169–185. 10.1007/BF00364149.

40. Du Lac, S., Raymond, J.L., Sejnowski, T.J., and Lisberger, S.G. (1995). Learning and memory in the vestibulo-ocular reflex. Annu. Rev. Neurosci. 18, 409–441.

41. Raymond, J.L., and Lisberger, S.G. (1998). Neural Learning Rules for the Vestibulo-Ocular Reflex. J. Neurosci. 18, 9112 LP – 9129. 10.1523/JNEUROSCI.18-21-09112.1998.

42. Carey, M.R., Medina, J.F., and Lisberger, S.G. (2005). Instructive signals for motor learning from visual cortical area MT. Nat. Neurosci. 8, 813–819. 10.1038/nn1470.

43. Shin, S.-L., Zhao, G.Q., and Raymond, J.L. (2014). Signals and Learning Rules Guiding Oculomotor Plasticity. J. Neurosci. 34, 10635 LP – 10644. 10.1523/JNEUROSCI.4510-12.2014.

44. Albert, S.T., and Shadmehr, R. (2016). The Neural Feedback Response to Error As a Teaching Signal for the Motor Learning System. J. Neurosci. 36, 4832 LP – 4845. 10.1523/JNEUROSCI.0159-16.2016.

45. Pugh, J.R., and Raman, I.M. (2009). Nothing can be coincidence: Synaptic inhibition and plasticity in the cerebellar nuclei. Trends Neurosci. 32, 170–177. 10.1016/j.tins.2008.12.001.

46. McElvain, L.E., Bagnall, M.W., Sakatos, A., and du Lac, S. (2010). Bidirectional plasticity gated by hyperpolarization controls the gain of postsynaptic firing responses at central vestibular nerve synapses. Neuron 68, 763–775. 10.1016/j.neuron.2010.09.025.

47. Lee, K.H., Mathews, P.J., Reeves, A.M.B., Choe, K.Y., Jami, S.A., Serrano, R.E., and Otis, T.S. (2015). Circuit Mechanisms Underlying Motor Memory Formation in the Cerebellum. Neuron 86, 529–540. 10.1016/j.neuron.2015.03.010.

48. Bonnan, A., Rowan, M.M.J., Baker, C.A., Bolton, M.M., and Christie, J.M. (2021). Autonomous Purkinje cell activation instructs bidirectional motor learning through evoked dendritic calcium signaling. Nat. Commun. 12, 2153. 10.1038/s41467-021-22405-8.

49. McCormick, D.A., Steinmetz, J.E., and Thompson, R.F. (1985). Lesions of the inferior olivary complex cause extinction of the classically conditioned eyeblink response. Brain Res. 359, 120–130. 10.1016/0006-8993(85)91419-2.

50. Luebke, A.E., and Robinson, D.A. (1994). Gain changes of the cat’s vestibulo-ocular reflex after flocculus deactivation. Exp. brain Res. 98, 379–390. 10.1007/BF00233976.

51. Zucca, R., Rasmussen, A., and Bengtsson, F. (2016). Climbing Fiber Regulation of Spontaneous Purkinje Cell Activity and Cerebellum-Dependent Blink Responses(1,2,3). eNeuro 3. 10.1523/ENEURO.0067-15.2015.

52. Lang, E.J., Apps, R., Bengtsson, F., Cerminara, N.L., De Zeeuw, C.I., Ebner, T.J., Heck, D.H., Jaeger, D., Jörntell, H., Kawato, M., et al. (2017). The Roles of the Olivocerebellar Pathway in Motor Learning and Motor Control. A Consensus Paper. The Cerebellum 16, 230–252. 10.1007/s12311-016-0787-8.

53. Albergaria, C., Silva, N.T., Pritchett, D.L., and Carey, M.R. (2018). Locomotor activity modulates associative learning in mouse cerebellum. Nat. Neurosci. 21, 725–735. 10.1038/s41593-018-0129-x.

54. Albergaria, C., Silva, N.T., Darmohray, D.M., and Carey, M.R. (2020). Cannabinoids modulate associative cerebellar learning via alterations in behavioral state. Elife 9, e61821. 10.7554/eLife.61821.

55. Gauck, V., and Jaeger, D. (2000). The Control of Rate and Timing of Spikes in the Deep Cerebellar Nuclei by Inhibition. J. Neurosci. 20, 3006 LP – 3016. 10.1523/JNEUROSCI.20-08-03006.2000.

56. Heiney, S.A., Kim, J., Augustine, G.J., and Medina, J.F. (2014). Precise Control of Movement Kinematics by Optogenetic Inhibition of Purkinje Cell Activity. J. Neurosci. 34, 2321 LP – 2330. 10.1523/JNEUROSCI.4547-13.2014.

57. Mostofi, A., Holtzman, T., Grout, A.S., Yeo, C.H., and Edgley, S.A. (2010). Electrophysiological Localization of Eyeblink-Related Microzones in Rabbit Cerebellar Cortex. J. Neurosci. 30, 8920 LP – 8934. 10.1523/JNEUROSCI.6117-09.2010.

58. Steinmetz, A.B., and Freeman, J.H. (2014). Localization of the cerebellar cortical zone mediating acquisition of eyeblink conditioning in rats. Neurobiol. Learn. Mem. 114, 148–154. 10.1016/j.nlm.2014.06.003.

59. Van Der Giessen, R.S., Koekkoek, S.K., van Dorp, S., De Gruijl, J.R., Cupido, A., Khosrovani, S., Dortland, B., Wellershaus, K., Degen, J., Deuchars, J., et al. (2008). Role of Olivary Electrical Coupling in Cerebellar Motor Learning. Neuron 58, 599–612. 10.1016/j.neuron.2008.03.016.

60. Hioki, H., Fujiyama, F., Taki, K., Tomioka, R., Furuta, T., Tamamaki, N., and Kaneko, T. (2003). Differential distribution of vesicular glutamate transporters in the rat cerebellar cortex. Neuroscience 117, 1–6. 10.1016/s0306-4522(02)00943-0.

61. Perrett, S.P., Ruiz, B.P., and Mauk, M.D. (1993). Cerebellar cortex lesions disrupt learning-dependent timing of conditioned eyelid responses. J. Neurosci. 13, 1708–1718. 10.1523/JNEUROSCI.13-04-01708.1993.

62. Chettih, S.N., McDougle, S.D., Ruffolo, L.I., and Medina, J.F. (2011). Adaptive timing of motor output in the mouse: the role of movement oscillations in eyelid conditioning. Front. Integr. Neurosci. 5, 72. 10.3389/fnint.2011.00072.

63. Tsubota, T., Ohashi, Y., Tamura, K., Sato, A., and Miyashita, Y. (2011). Optogenetic Manipulation of Cerebellar Purkinje Cell Activity In Vivo. PLoS One 6, e22400.

64. Canto, C.B., Witter, L., and De Zeeuw, C.I. (2016). Whole-Cell Properties of Cerebellar Nuclei Neurons In Vivo. PLoS One 11, e0165887.

65. El-Shamayleh, Y., Kojima, Y., Soetedjo, R., and Horwitz, G.D. (2017). Selective Optogenetic Control of Purkinje Cells in Monkey Cerebellum. Neuron 95, 51–62.e4. 10.1016/j.neuron.2017.06.002.

66. Menardy, F., Varani, A.P., Combes, A., Léna, C., and Popa, D. (2019). Functional Alteration of Cerebello-Cerebral Coupling in an Experimental Mouse Model of Parkinson’s Disease. Cereb. Cortex 29, 1752–1766. 10.1093/cercor/bhy346.

67. Jackman, S.L., Chen, C.H., Offermann, H.L., Drew, I.R., Harrison, B.M., Bowman, A.M., Flick, K.M., Flaquer, I., and Regehr, W.G. (2020). Cerebellar Purkinje cell activity modulates aggressive behavior. Elife 9, e53229. 10.7554/eLife.53229.

68. Bina, L., Romano, V., Hoogland, T.M., Bosman, L.W.J., and De Zeeuw, C.I. (2021). Purkinje cells translate subjective salience into readiness to act and choice performance. Cell Rep. 37, 110116. 10.1016/j.celrep.2021.110116.

69. Guo, C., Witter, L., Rudolph, S., Elliott, H.L., Ennis, K.A., and Regehr, W.G. (2016). Purkinje Cells Directly Inhibit Granule Cells in Specialized Regions of the Cerebellar Cortex. Neuron 91, 1330–1341. 10.1016/j.neuron.2016.08.011.

70. Guo, C., Rudolph, S., Neuwirth, M.E., and Regehr, W.G. (2021). Purkinje cell outputs selectively inhibit a subset of unipolar brush cells in the input layer of the cerebellar cortex. Elife 10. 10.7554/eLife.68802.

71. Orduz, D., and Llano, I. (2007). Recurrent axon collaterals underlie facilitating synapses between cerebellar Purkinje cells. Proc. Natl. Acad. Sci. U. S. A. 104, 17831–17836. 10.1073/pnas.0707489104.

72. Ankri, L., Husson, Z., Pietrajtis, K., Proville, R., Léna, C., Yarom, Y., Dieudonné, S., and Uusisaari, M.Y. (2015). A novel inhibitory nucleo-cortical circuit controls cerebellar Golgi cell activity. Elife 4, e06262. 10.7554/eLife.06262.

73. Houck, B.D., and Person, A.L. (2015). Cerebellar Premotor Output Neurons Collateralize to Innervate the Cerebellar Cortex. J. Comp. Neurol. 523, 2254–2271. 10.1002/cne.23787.

74. Garcia, K.S., Steele, P.M., and Mauk, M.D. (1999). Cerebellar Cortex Lesions Prevent Acquisition of Conditioned Eyelid Responses. J. Neurosci. 19, 10940 LP – 10947. 10.1523/JNEUROSCI.19-24-10940.1999.

75. Herman, A.M., Huang, L., Murphey, D.K., Garcia, I., and Arenkiel, B.R. (2014). Cell type-specific and time-dependent light exposure contribute to silencing in neurons expressing Channelrhodopsin-2. Elife 3, e01481. 10.7554/eLife.01481.

76. Chaumont, J., Guyon, N., Valera, A.M., Dugué, G.P., Popa, D., Marcaggi, P., Gautheron, V., Reibel-Foisset, S., Dieudonné, S., Stephan, A., et al. (2013). Clusters of cerebellar Purkinje cells control their afferent climbing fiber discharge. Proc. Natl. Acad. Sci. 110, 16223–16228. 10.1073/pnas.1302310110.

77. Grangeray-Vilmint, A., Valera, A.M., Kumar, A., and Isope, P. (2018). Short-Term Plasticity Combines with Excitation–Inhibition Balance to Expand Cerebellar Purkinje Cell Dynamic Range. J. Neurosci. 38, 5153 LP – 5167. 10.1523/JNEUROSCI.3270-17.2018.

78. Chuong, A.S., Miri, M.L., Busskamp, V., Matthews, G.A.C., Acker, L.C., Sørensen, A.T., Young, A., Klapoetke, N.C., Henninger, M.A., Kodandaramaiah, S.B., et al. (2014). Noninvasive optical inhibition with a red-shifted microbial rhodopsin. Nat. Neurosci. 17, 1123–1129. 10.1038/nn.3752.

79. Aschauer, D.F., Kreuz, S., and Rumpel, S. (2013). Analysis of Transduction Efficiency, Tropism and Axonal Transport of AAV Serotypes 1, 2, 5, 6, 8 and 9 in the Mouse Brain. PLoS One 8, e76310.

80. Titley, H.K., Kislin, M., Simmons, D.H., Wang, S.S.-H., and Hansel, C. (2019). Complex spike clusters and false-positive rejection in a cerebellar supervised learning rule. J. Physiol. 597, 4387–4406. 10.1113/JP278502.

81. Suriano, C.M., Verpeut, J.L., Kumar, N., Ma, J., Jung, C., and Boulanger, L.M. (2021). Adeno-associated virus (AAV) reduces cortical dendritic complexity in a TLR9-dependent manner. bioRxiv, 2021.09.28.462148. 10.1101/2021.09.28.462148.

82. Zimmermann, D., Zhou, A., Kiesel, M., Feldbauer, K., Terpitz, U., Haase, W., Schneider-Hohendorf, T., Bamberg, E., and Sukhorukov, V.L. (2008). Effects on capacitance by overexpression of membrane proteins. Biochem. Biophys. Res. Commun. 369, 1022– 1026. 10.1016/j.bbrc.2008.02.153.

83. Lin, J.Y. (2012). Chapter 2 - Optogenetic excitation of neurons with channelrhodopsins: Light instrumentation, expression systems, and channelrhodopsin variants. In Optogenetics: Tools for Controlling and Monitoring Neuronal Activity, T. Knöpfel and E. S. B. T.-P. in B. R. Boyden, eds. (Elsevier), pp. 29–47. 10.1016/B978-0-444-59426-6.00002-1.

84. Miyashita, T., Shao, Y., Chung, J., Pourzia, O., and Feldman, D. (2013). Long-term channelrhodopsin-2 (ChR2) expression can induce abnormal axonal morphology and targeting in cerebral cortex. Front. Neural Circuits 7.

85. Liu, M., Sharma, A.K., Shaevitz, J.W., and Leifer, A.M. (2018). Temporal processing and context dependency in Caenorhabditis elegans response to mechanosensation. Elife 7, e36419. 10.7554/eLife.36419.

86. Garden, D.L.F., Oostland, M., Jelitai, M., Rinaldi, A., Duguid, I., and Nolan, M.F. (2018). Inferior Olive HCN1 Channels Coordinate Synaptic Integration and Complex Spike Timing. Cell Rep. 22, 1722–1733. 10.1016/j.celrep.2018.01.069.

87. Lefler, Y., Yarom, Y., and Uusisaari, M.Y. (2014). Cerebellar inhibitory input to the inferior olive decreases electrical coupling and blocks subthreshold oscillations. Neuron 81, 1389–1400. 10.1016/j.neuron.2014.02.032.

88. Bazzigaluppi, P., Isenia, S.C., Haasdijk, E.D., Elgersma, Y., De Zeeuw, C.I., van der Giessen, R.S., and de Jeu, M.T.G. (2017). Modulation of Murine Olivary Connexin 36 Gap Junctions by PKA and CaMKII. Front. Cell. Neurosci. 11.

89. Powell, K., Mathy, A., Duguid, I., and Häusser, M. (2015). Synaptic representation of locomotion in single cerebellar granule cells. Elife 4, e07290. 10.7554/eLife.07290.

90. Ishikawa, T., Shimuta, M., and Häusser, M. (2015). Multimodal sensory integration in single cerebellar granule cells in vivo. Elife 4, e12916. 10.7554/eLife.12916.

91. Wang, Y.T., and Linden, D.J. (2000). Expression of Cerebellar Long-Term Depression Requires Postsynaptic Clathrin-Mediated Endocytosis. Neuron 25, 635–647. 10.1016/S0896-6273(00)81066-1.

92. Roome, C.J., and Kuhn, B. (2018). Simultaneous dendritic voltage and calcium imaging and somatic recording from Purkinje neurons in awake mice. Nat. Commun. 9, 3388. 10.1038/s41467-018-05900-3.

93. Roome, C.J., and Kuhn, B. (2020). Dendritic coincidence detection in Purkinje neurons of awake mice. Elife 9, e59619. 10.7554/eLife.59619.

94. Hartell (1996). Strong Activation of Parallel Fibers Produces Localized Calcium Transients and a Form of LTD That Spreads to Distant Synapses. Neuron 16, 601–610. 10.1016/S0896-6273(00)80079-3.

95. Hartell (2002). Parallel fiber plasticity. The Cerebellum 1, 3–18. 10.1080/147342202753203041.

96. Herzfeld, D.J., Kojima, Y., Soetedjo, R., and Shadmehr, R. (2015). Encoding of action by the Purkinje cells of the cerebellum. Nature 526, 439–442. 10.1038/nature15693.

97. Herzfeld, D.J., Kojima, Y., Soetedjo, R., and Shadmehr, R. (2018). Encoding of error and learning to correct that error by the Purkinje cells of the cerebellum. Nat. Neurosci. 21, 736–743. 10.1038/s41593-018-0136-y.

98. Darmohray, D.M., Jacobs, J.R., Marques, H.G., and Carey, M.R. (2019). Spatial and Temporal Locomotor Learning in Mouse Cerebellum. Neuron 102, 217–231.e4. 10.1016/j.neuron.2019.01.038.

99. Broersen, R., Albergaria, C., Carulli, D., Carey, M.R., Canto, C.B., and Zeeuw, C.I. De (2023). Synaptic mechanisms for associative learning in the cerebellar nuclei. bioRxiv, 2022.10.28.514163. 10.1101/2022.10.28.514163.

100. Kassardjian, C.D., Tan, Y.-F., Chung, J.-Y.J., Heskin, R., Peterson, M.J., and Broussard, D.M. (2005). The Site of a Motor Memory Shifts with Consolidation. J. Neurosci. 25, 7979 LP – 7985. 10.1523/JNEUROSCI.2215-05.2005.

101. Yang, Y., and Lisberger, S.G. (2010). Learning on Multiple Timescales in Smooth Pursuit Eye Movements. J. Neurophysiol. 104, 2850–2862. 10.1152/jn.00761.2010.

102. Medina, J.F., Garcia, K.S., and Mauk, M.D. (2001). A Mechanism for Savings in the Cerebellum. J. Neurosci. 21, 4081 LP – 4089. 10.1523/JNEUROSCI.21-11-04081.2001.

103. Lisberger, S.G. (1994). Neural basis for motor learning in the vestibuloocular reflex of primates. III. Computational and behavioral analysis of the sites of learning. J. Neurophysiol. 72, 974–998. 10.1152/jn.1994.72.2.974.

104. Madisen, L., Mao, T., Koch, H., Zhuo, J., Berenyi, A., Fujisawa, S., Hsu, Y.-W.A., Garcia, A.J. 3rd, Gu, X., Zanella, S., et al. (2012). A toolbox of Cre-dependent optogenetic transgenic mice for light-induced activation and silencing. Nat. Neurosci. 15, 793–802. 10.1038/nn.3078.

105. Barski, J.J., Dethleffsen, K., and Meyer, M. (2000). Cre recombinase expression in cerebellar Purkinje cells. Genesis 28, 93–98.

106. Fünfschilling, U., and Reichardt, L.F. (2002). Cre-mediated recombination in rhombic lip derivatives. genesis 33, 160–169. 10.1002/gene.10104.

107. Carey, M.R., Myoga, M.H., McDaniels, K.R., Marsicano, G., Lutz, B., Mackie, K., and Regehr, W.G. (2010). Presynaptic CB1 Receptors Regulate Synaptic Plasticity at Cerebellar Parallel Fiber Synapses. J. Neurophysiol. 105, 958–963. 10.1152/jn.00980.2010.

108. Fremeau, R.T., Troyer, M.D., Pahner, I., Nygaard, G.O., Tran, C.H., Reimer, R.J., Bellocchio, E.E., Fortin, D., Storm-Mathisen, J., and Edwards, R.H. (2001). The Expression of Vesicular Glutamate Transporters Defines Two Classes of Excitatory Synapse. Neuron 31, 247–260. 10.1016/S0896-6273(01)00344-0.

109. Borgius, L., Restrepo, C.E., Leao, R.N., Saleh, N., and Kiehn, O. (2010). A transgenic mouse line for molecular genetic analysis of excitatory glutamatergic neurons. Mol. Cell. Neurosci. 45, 245–257. 10.1016/j.mcn.2010.06.016.

110. Lee, J.H., Durand, R., Gradinaru, V., Zhang, F., Goshen, I., Kim, D.-S., Fenno, L.E., Ramakrishnan, C., and Deisseroth, K. (2010). Global and local fMRI signals driven by neurons defined optogenetically by type and wiring. Nature 465, 788–792. 10.1038/nature09108.

111. De Zeeuw, C.I., Wentzel, P., and Mugnaini, E. (1993). Fine structure of the dorsal cap of the inferior olive and its GAB aergic and non-Gabaergic input from the nucleus prepositus hypoglossi in rat and rabbit. J. Comp. Neurol. 327, 63–82. 10.1002/cne.903270106.

112. Arenkiel, B.R., Peca, J., Davison, I.G., Feliciano, C., Deisseroth, K., Augustine, G.J., Ehlers, M.D., and Feng, G. (2007). In Vivo Light-Induced Activation of Neural Circuitry in Transgenic Mice Expressing Channelrhodopsin-2. Neuron 54, 205–218. 10.1016/j.neuron.2007.03.005.

113. Markanday, A., Bellet, J., Bellet, M.E., Inoue, J., Hafed, Z.M., and Thier, P. (2020). Using deep neural networks to detect complex spikes of cerebellar Purkinje cells. J. Neurophysiol. 123, 2217–2234. 10.1152/jn.00754.2019.

114. Wang, X., Liu, Z., Angelov, M., Feng, Z., Li, X., Li, A., Yang, Y., Gong, H., and Gao, Z. (2023). Excitatory nucleo-olivary pathway shapes cerebellar outputs for motor control. Nat. Neurosci. 26, 1394–1406. 10.1038/s41593-023-01387-4.

